# Spatial ciliary signaling regulates the dorsal/ventral regionalization of human brain organoids

**DOI:** 10.1101/2024.07.18.604098

**Authors:** Issei S. Shimada, Akari Goto, Yutaka Hashimoto, Hiroshi Takase, Masayuki Itoh, Yoichi Kato

## Abstract

Regionalization of the brain is a fundamental question in human developmental biology. Primary cilia are known for a critical organelle for dorsal/ventral fate of brain formation in mice, but little is known about how signaling in the primary cilia regulate regionalization of the human brain. Here, we found that signaling in the primary cilia function in regionalization of the brain using brain organoids derived from human induced pluripotent stem (iPS) cells. Deletion of a ciliary GTPase, *ARL13B*, induced partially ventralized neural stem cells in the dorsal cortical organoids, despite using a guided dorsal cortical organoid differentiation protocol. Mechanistically, *ARL13B* knockout (KO) neural stem cells decreased ciliary localization of GPR161, a negative regulator of SHH signaling in primary cilia and increased SONIC HEDGEHOG (SHH) signaling. *GPR161* deletion also induced ventralized neural stem cells in the dorsal cortical organoids, despite using the guided differentiation protocol. *GPR161* deletion increased SHH signaling mediated by decreased GLI3 repressor formation. Pharmacological treatment to increase cAMP levels rescued GLI3 repressor formation and the differentiation of dorsal neural stem cells in *GPR161* KO brain organoids. Importantly, elevating the amount of ciliary cAMP by optogenetics restored the generation of dorsal neural stem cells in *GPR161* KO brain organoids. These data indicate that spatial ciliary signaling, the ARL13B-GPR161-cAMP axis in primary cilia, is a fundamental regulator of the dorsal/ventral regionalization of the human brain.

## Introduction

Brain regionalization is a critical process for human brain development. Although the general characteristics of brain function and development have been studied in various animal models, it remains elusive what determines our brain architecture ^1–3^. Neural stem cells, also known as neural progenitor cells or radial glia, hold the key to this enigma ^4^. Neural stem cells are the most primitive cell types in brain development and generate neurons and other cell types during brain development ^3,5–7^. Mutations of essential genes in neural stem cell maintenance result in decreased self-renewal and multipotent capacities ^8^. Impaired function of neural stem cells results in compromised neuronal and glial differentiation, leading to the failure of neural network formation ^9^. Indeed, dysfunction of neural stem cells leads various diseases, such as microcephaly, autism, fragile X syndrome and neural tube defects ^10–15^. Therefore, the analysis of neural stem cells can lead to a better understanding of the mechanisms that govern human brain regionalization.

Primary cilium is an antenna-like organelle with various biological functions, including cell signaling in cancer and development ^16–18^. To function as an antenna for extracellular signaling, primary cilia contain ion channels, more than 30 G-protein coupled receptors (GPCRs), adenylyl cyclases (3, 5 and 6), PKA holoenzymes and their substrates ^19–21^. Primary cilia function as post-translational regulators of various transcription factors, synapse recipients and specialized cAMP switch, enhancing the importance of primary cilia in our body ^16,22,23^. Importantly, most (if not all) neural stem cells have primary cilia which protrudes into the ventricle ^24^. Ciliary dysfunction leads to various brain malformation such as neural tube defects, Joubert syndrome and Dandy-Walker syndrome, which are commonly classified as ciliopathies ^25^. However, it is not well understood how disorders of cilia give rise to the human brain malformation.

Sonic hedgehog (SHH) signaling is one of the major signal transduction pathways involved in the function of primary cilia. SHH functions as a cytokine, and is critical for both developmental biology and cancer biology ^26,27^. SHH interacts with its receptor, patched 1 (PTCH1), on the ciliary membrane, which induces delocalization of PTCH1 from the ciliary membrane and localization of SHH signaling activator, smoothened (SMO) into primary cilia ^28^. SMO utilized cholesterol in the membrane of primary cilia and activates SHH signaling by modifying the phosphorylation status of GLI2 transcription factor (GLI-Activator) ^29,30^. SHH signaling is basically suppressed by two distinct mechanisms. One mechanism is that PTCH1 hinders SMO localization into cilia and prevents the formation of GLI-Activator. Another mechanism is that an orphan G protein-coupled receptor 161 (GPR161) constitutively activates cAMP/PKA signaling, resulting in the phosphorylation of GLI3 and the cleavage of full-length GLI3 into the repressor form (GLI-Repressor) ^13,31,32^. This process leads to transcriptional inhibition of the SHH signaling pathway ^33^. These precise spatial, post-translational and transcriptional mechanisms determine the strength of the SHH signaling in our body.

SHH signaling through primary cilia plays a critical role in telencephalic patterning^34–37^. As development progresses, the activation of SHH signaling causes expansion of cortical surface area by inducing an increased number of neural stem cells, also known as apical radial glia at this stage, and basal radial glia ^12,38,39^. During embryonic stage, SHH signaling is deeply involved in neural tube development ^40,41^.

SHH is secreted from the notochord and floor plate of the neural tube, and induces ventral identity in neural stem cells ^42^. On the other hand, WNT and BMP are secreted from the dorsal region of the neural tube, and induce dorsal identity in neural stem cells ^43–45^. The disruption of SHH signaling induces altered dorsal/ventral fate in neural tube formation ^46,47^. Upregulated SHH signaling induces ventralization of neural tube, while downregulated SHH signaling induces dorsalization in mice ^41,48^.

Recent studies have utilized the human brain to elucidate the developmental mechanisms. However, it is not easy to obtain and analyze human brain ^6,49^. Brain organoids are the *in vitro* model of human brain generated from pluripotent stem cells ^15,50,51^. Various regions of the brain have been generated, including the cortex, thalamus, hippocampus, cerebellum, spinal cord and hypothalamus ^15,52–56^. Previous studies showed the existence of primary cilia in the cortical brain organoids ^51,57,58^; however, little is known how primary cilia is critical in human brain regionalization and whether signaling in the primary cilia is sufficient for regionalization of the brain. Here, we characterize the ciliary signaling axis that regulates the dorsal/ventral fate determination of brain structure using brain organoid models.

## Results

### Primary cilia point towards the ventricle in brain organoid models

Neural stem cells possessing primary cilia are observed in both human, mice and brain organoid models, but the functions of primary cilia in neural stem cell regulation is not well known ^51,57,58^. To examine the role of primary cilia in human neural stem cells, we generated induced pluripotent stem (iPS) cell-derived dorsal cortical organoids using a guided differentiation protocol ^59^ (Figure 1A). After 1 month of culture, the dorsal cortical organoid contained multiple ventricular zones (Figure 1B). The ventricular zones were composed of BLBP-positive neural stem cells surrounded by beta-tubulin III-positive neurons (Figure 1C). As similar to the mouse brain, phosphorylated histone H3 (pH3)-positive mitosis occurred at the inner surface of PAX6-positive ventricular zones (Figure 1D). Electron microscopy analysis also revealed accumulation of mitotic cells around the ventricular zones (Figure 1E). ARL13B-positive primary cilia were densely packed in the ventricles of dorsal cortical organoids (Figure 1F). We next performed scanning electron microscopy analysis to visualize the localization and the morphology of primary cilia. We observed primary cilia pointed toward the ventricles (Figure 1G-I). In transmission electron microscopy analysis, we clearly visualized that each basal foot of neural stem cells contained one primary cilium (Figure 1J, K). We did not observe any neural stem cells containing more than two primary cilia. Each basal foot of neural stem cells contained one mother centriole and one daughter centriole (Figure 1L). The electron microscope image of the cross section of primary cilia showed a 9+0 axoneme structure, which strongly supports that neural stem cells have primary cilia in the dorsal cortical organoids. Thus, neural stem cells in dorsal cortical organoids contained primary cilia pointing towards the ventricles, similar to the human fetus brain.

**Figure 1.**
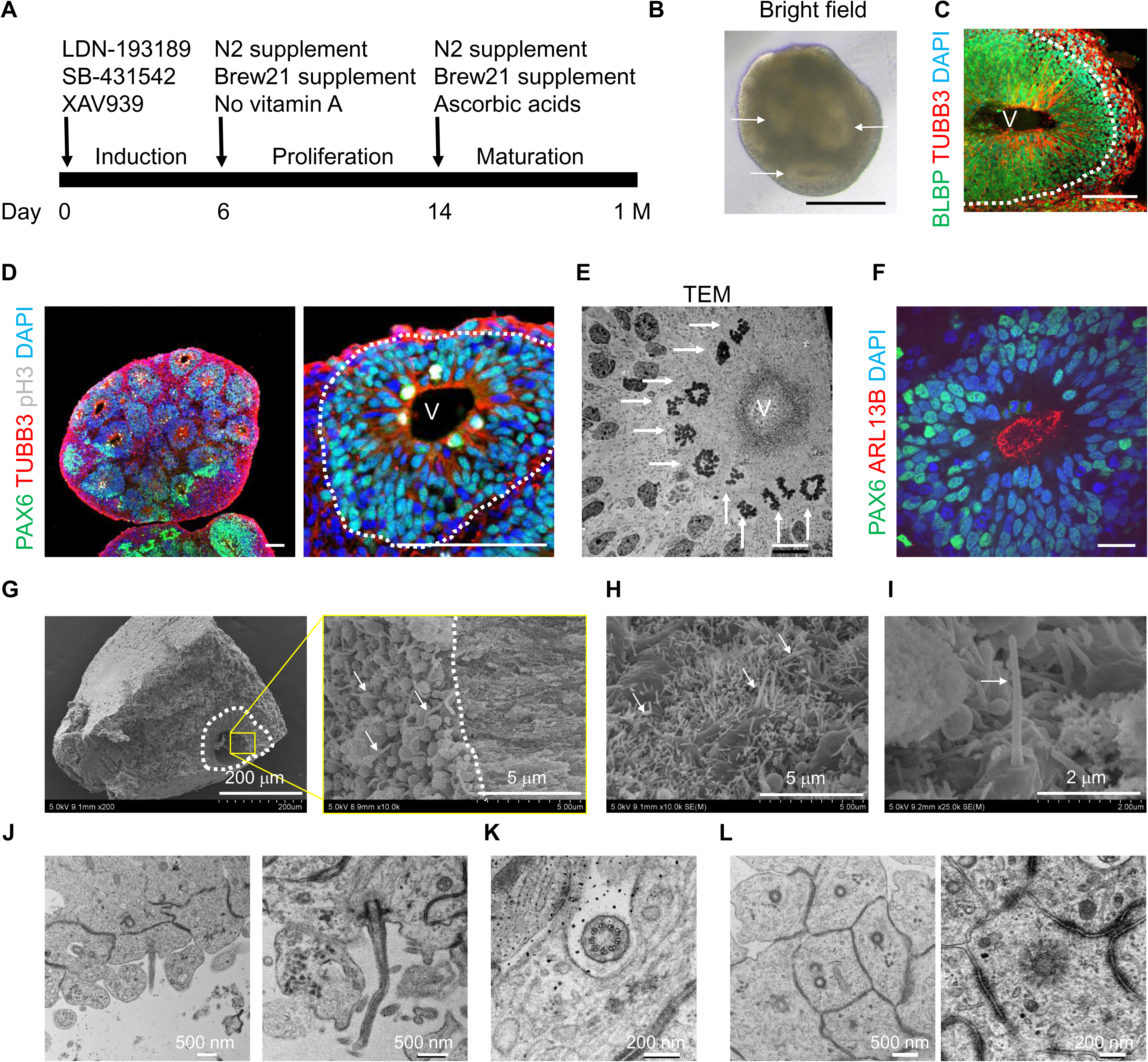
Human neural stem cells had primary cilia in the dorsal cortical organoids. (**A**) A scheme of dorsal cortical organoid generation from iPS cells. M indicates a month unit. (**B**) A bright-field view of a human dorsal cortical organoid. Arrows indicate the ventricles. (**C**) BLBP-positive neural stem cells and beta-tubulin III(TUBB3)-positive neurons were observed in an inside-outside manner in the dorsal cortical organoid. White dotted line indicates the junction between ventricular zone and cortical plate. V, ventricle. (**D**) pH3-positive mitotic cells were observed in the inner layer of PAX6-positive neural stem cells enriched in the ventricular zone. Beta-tubulin III (TUBB3)-positive neurons were observed around the ventricular zone. (**E**) Transmission electron microscope (TEM) analysis revealed the existence of mitotic cells in the inner layer of the ventricular zone. Arrows indicate mitotic cells. (**F**) ARL13B-positive primary cilia were observed inside of the ventricular zone enriched with PAX6-positive neural stem cells. (**G**–**I**) Scanning electron microscope (SEM) analysis revealed a primary cilia-enriched region on the inner side of the ventricular zone of human dorsal cortical organoids. White dotted lines indicate ventricular zones. Arrows indicate representative primary cilia. (**J**) TEM analysis revealed the existence of one primary cilium per neural stem cell in the ventricular zone. (**K**) TEM analysis revealed that the primary cilium in the ventricular zone has a 9+0 structure. (**L**) TEM analysis revealed the existence of basal bodies at the apical end of neural stem cells in the ventricular zone. Scale bars are (**B**) 500 µm, (**C**) 100 µm, (**D**) 100 µm, (**E**) 10 µm, (**F**) 20 µm, (**G**, left) 200 µm, (**G**, right) 5 µm, (**H**) 5 µm, (**I**) 2 µm, (**J**) 500 nm, (**K**) 200 nm, (**L**, left) 500 nm, and (**L**, right) 200 nm.

### Deletion of *ARL13B* induces disrupted morphology of primary cilia in dorsal neural stem cells

ARL13B is a small GTPase enriched in primary cilia, and contributes to cilia morphology and ciliary signaling ^60–62^. Deletion of *ARL13B* alters membrane components, receptor localization, and protein trafficking in primary cilia, therefore resulting in the dysfunction of ciliary signaling. To examine how ciliary signaling is critical for neural stem cells in dorsal cortical organoids, we generated *ARL13B* knockout (KO) iPS cells using CRISPR/Cas9 system (Figure 2A, B, Supplemental Figure 1A). Deletion of *ARL13B* in iPS cells was confirmed by immunocytochemistry (Figure 2C, D). Deletion of *ARL13B* did not affect the cell growth (Figure 2E) and the stem cell marker expression (Figure 2F) of iPS cells compared to control iPS cells. These data indicate that ARL13B may be dispensable for iPS cell maintenance. To examine the function of ciliary signaling in neural stem cells, we generated dorsal cortical organoids derived from *ARL13B* KO iPS cells using a guided differentiation protocol. We first confirmed that ARL13B protein was absent in *ARL13B* KO dorsal cortical organoids at 1 month of culture (Figure 2G, H). Deletion of *ARL13B* did not disrupt the generation of PAX6-positive neural stem cells in the dorsal cortical organoids as compare to control organoids (Figure 2I, J).

**Figure 2.**
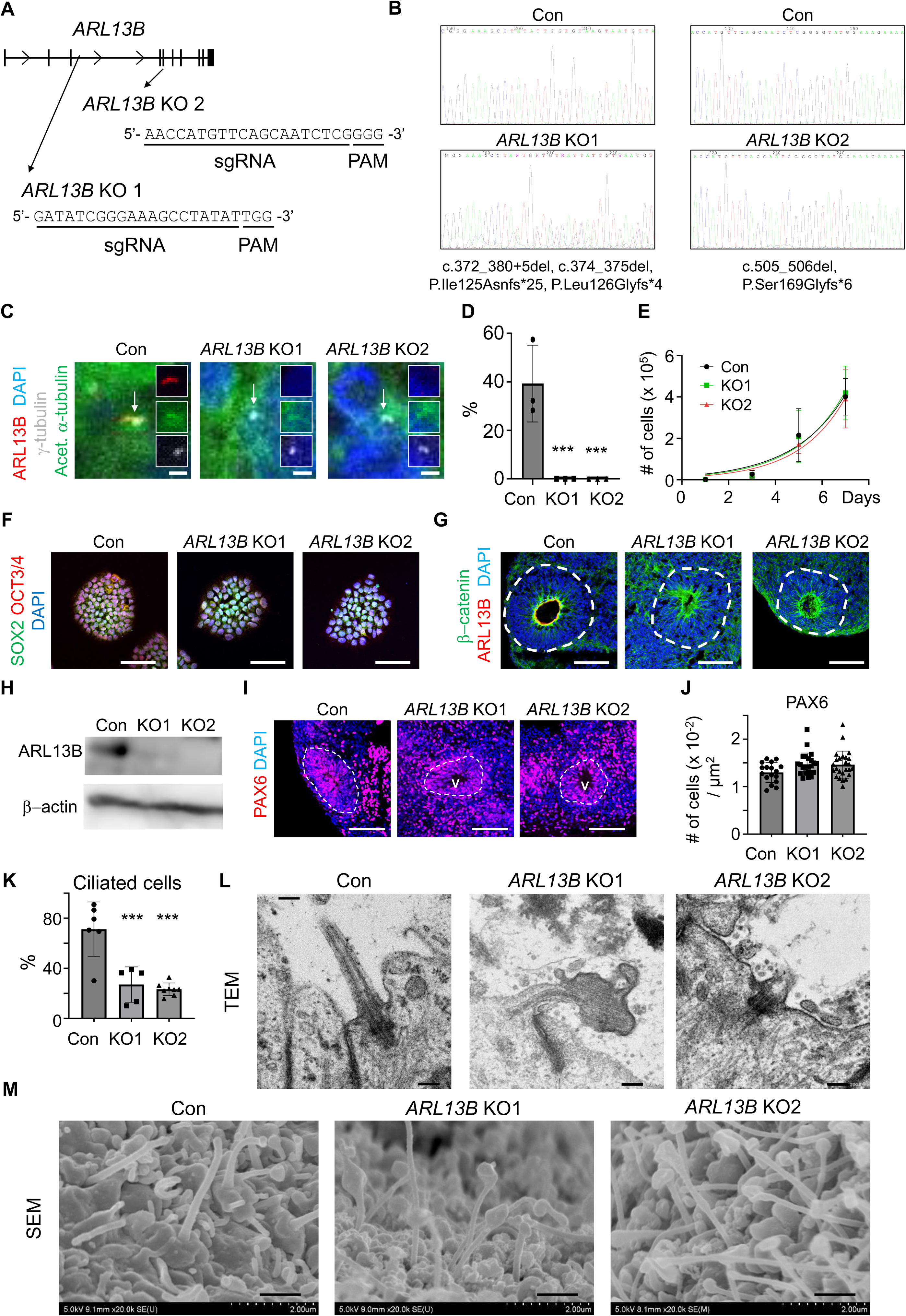
Deletion of *ARL13B* induced disrupted morphology of primary cilia in neural stem cells in human dorsal cortical organoids. (**A**) A scheme of *ARL13B* gene with CRISPR/Cas9 target sequences. (**B**) Sanger sequences of the CRISPR/Cas9-targeted loci in control (Con), *ARL13B* KO1 and *ARL13B* KO2 iPS cells (#51). (**C**) ARL13B-positive primary cilia were shown in control, but not in *ARL13B* KO1 and *ARL13B* KO2 iPS cells. gamma-tubulin and acetylated alpha-tubulin indicate basal bodies and primary cilia, respectively. Arrows indicate basal bodies. (**D**) Quantification of ARL13B-positive primary cilia in (**C**). N = 3 technical replicates. (**E**) Cell growth curves of control, *ARL13B* KO1 and *ARL13B* KO2 iPS cells. N = 3 technical replicates. (**F**) SOX2 and OCT3/4 expression in control, *ARL13B* KO1 and *ARL13B* KO2 iPS cells. (**G**) ARL13B-positive primary cilia were shown in the ventricular zone of control, but not of *ARL13B* KO1 and *ARL13B* KO2 dorsal cortical organoids. White dotted lines indicate ventricular zones. Beta-catenin-positive cells represent neural stem cells. (**H**) Western blot analysis revealed that ARL13B expression was absent in *ARL13B* KO1 and *ARL13B* KO2 dorsal cortical organoids. β-actin is a loading control. (**I**) PAX6-positive neural stem cells were shown in the ventricular zone of control, *ARL13B* KO1 and *ARL13B* KO2 dorsal cortical organoids. V, ventricle. (**J**) The number of PAX6-positive neural stem cells in the ventricular zone of control, *ARL13B* KO1 and *ARL13B* KO2 cortical organoids. N = 11–15 ventricular zones of dorsal cortical organoids. (**K**) Comparison of the percentage of ciliated cells using control, *ARL13B* KO1 and *ARL13B* KO2 neural stem cells in 2D culture. N= 5–8 region of interests in 2 independent experiments. *** p < 0.001 to control. (**L**) TEM analysis revealed the existence of bulged and shortened primary cilia in the ventricular zone of *ARL13B* KO1 and *ARL13B* KO2 dorsal cortical organoids. (**M**) SEM analysis revealed the existence of bulged and shortened primary cilia in the ventricular zone of *ARL13B* KO1 and *ARL13B* KO2 dorsal cortical organoids. Scale bars are (**C**) 2 µm, (**F**) 100 µm, (**G**) 100 µm, (**I**) 100 µm, (**L**) 200 nm, and (**M**) 10 µm.

To examine the effects of *ARL13B* deletion in primary cilia, we cultured iPS cell-derived neural stem cells in 2D condition and quantified the number and length of primary cilia. The number of ciliated cells were significantly decreased in *ARL13B* KO neural stem cells in the 2D culture (Figure 2K, Supplemental Figure 2A). The length of cilia was significantly shorter in *ARL13B* KO neural stem cells than in control cells (Supplemental Figure 2B). To analyze primary cilia in *ARL13B* KO dorsal cortical organoids, we performed an electron microscopy analysis. Primary cilia were easily visualized in the ventricles of dorsal cortical organoids (Figure 2L). In some cases, we identified primary cilia whose tips were bulged in *ARL13B* KO neural stem cells (Figure 2L, middle). Microtubule-like structure was accumulated at the tip of primary cilia in *ARL13B* KO neural stem cells, suggesting disrupted cargo trafficking in primary cilia. Scanning electron microscopy analysis revealed an enrichment of bulged primary cilia in the ventricle of *ARL13B* KO dorsal cortical organoids (Figure 2M). Previous studies showed that the disruption of IFT-A complex, which is required for the retrograde transport to the basal body, also induced bulged primary cilia, suggesting that *Arl13B* deletion induced disrupted IFT-A function in cilia ^63–65^. The data indicate that *ARL13B* deletion disrupted the morphology of primary cilia in neural stem cells of dorsal cortical organoids.

### *ARL13B* deletion activates the SHH signaling pathway and induces partial ventralization

ARL13B is a critical component of ciliary signaling and regulates the SHH signaling pathway in a context dependent manner ^62,66^. To examine the effects of disrupted primary cilia on neural stem cells, we isolated RNAs from neural stem cells in dorsal cortical organoids before the neural maturation period and examined the expression levels of SHH target genes by qRT-PCR. *ARL13B* deletion significantly upregulated the expression of *GLI1* but weakly downregulated those of *GLI2* and *PTCH1* in neural stem cells (Figure 3A). Since PTCH1 and GLI regulate the SHH signaling pathway through multiple feedback mechanisms in a context dependent manner ^67^, these data indicate that ARL13B contributes to the SHH signaling pathway via the regulation of PTCH1 and GLI. Since the SHH signaling pathway is critical for dorsal/ventral determination of neural stem cells during neural tube formation ^68^, we examined the effects of SHH signaling in dorsal cortical brain organoids. Almost all cells in the ventricular zone expressed SOX2, indicating that the cells in the ventricular zones were neural stem cells (Figure 3B, C). Interestingly, *ARL13B*-deleted iPS cells-derived dorsal cortical organoids contained NKX2.2-positive neural stem cells, indicating the appearance of ventral neural stem cells (Figure 3B, D). These data are surprising, as we employed the guided differentiation protocol to generate dorsal cortical brain organoids using dual SMAD inhibitors ^59^. We also observed an increased number of OLIG2-positive neural stem cells but not FOXA2-positive neural stem cells in *ARL13B* KO dorsal cortical organoids compared to control dorsal cortical organoids, indicating mild ventral induction by *ARL13B* deletion (Figure 3E–H). Interestingly, since cells in the ventricular zones expressed PAX6, a marker of dorsal neural stem cells (Figure 2I, J), the loss of *ARL13B* does not cause complete ventralization of the organoids.

**Figure 3.**
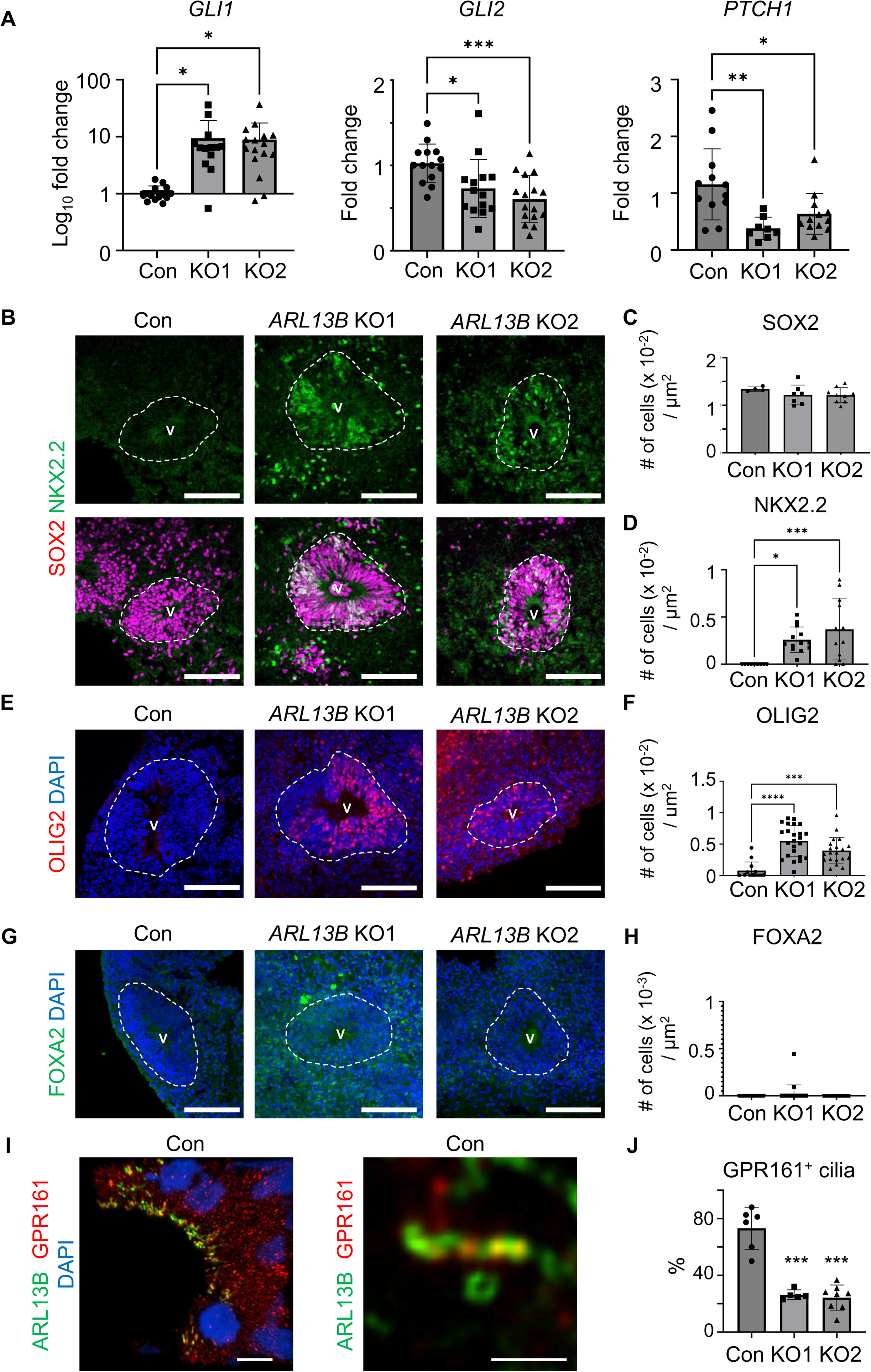
*ARL13B* deletion induced mild ventralization in dorsal cortical organoids. (**A**) qRT-PCR revealed modified expression patterns of SHH target genes in *ARL13B* KO1 and *ARL13B* KO2 dorsal cortical organoids during the proliferation period. Each dot indicates mRNAs derived from 10-30 brain organoids. N = 8–16 accumulation of brain organoids. (**B**–**D**) SOX2-positive pan neural stem cells and NKX2.2-positive ventralized neural stem cells were observed in *ARL13B* KO1 and *ARL13B* KO2 dorsal cortical organoids. White dotted lines indicate ventricular zones. V, ventricle. N = 4–13 ventricular zones of dorsal cortical organoids. (**E**, **F**) OLIG2-positive ventralized neural stem cells were observed in *ARL13B* KO1 and *ARL13B* KO2 brain organoids. N = 13–24 ventricular zones of dorsal cortical organoids. (**G, H**) FOX2A-positive ventralized neural stem cells were not observed in *ARL13B* KO1 and *ARL13B* KO2 dorsal cortical organoids. N = 12–15 ventricular zones of dorsal cortical organoids. (**I**) GPR161 localization in ARL13B-positive primary cilia in the ventricular zone of control dorsal cortical organoids. (**J**) Quantification of GPR161 positive cilia in control, *ARL13B* KO1 and *ARL13B* KO2 neural stem cells in 2D cutlure. N = 5–8 region of interests in 2 independent experiments. * indicates p < 0.05,** indicates p < 0.01 and *** indicates p < 0.001 to control. Scale bars are (**B**) 100 µm, (**E**) 100 µm, (**G**) 100 µm, (**I**, left) 5 µm and (**I**, right) 1 µm.

### GPR161 negatively regulates the SHH signaling pathway

GPR161 localizes to primary cilia and negatively regulates the SHH signaling pathway ^12,13,69,70^. Previous studies showed that deletion of *ARL13B* caused GPR161 mislocalization and impaired responsiveness to SHH signaling in primary cilia ^71,72^. We observed that GPR161 localized to primary cilia of neural stem cells of the dorsal cortical brain organoids (Figure 3I). The percentages of GPR161 positive cilia were significantly decreased in *ARL13B* KO neural stem cells in 2D culture system (Figure 3J, Supplementary Figure 2A). To examine the role of GPR161 in dorsal cortical brain organoids, we generated *GPR161* KO iPS cells using CRISPR/Cas9 system (Figure 4A, B, Supplemental Figure 1B). The deletion of *GPR161* did not affect the expression of stem cell markers in iPS cells(Figure 4C). The cell growth of *GPR161* KO iPS cells was similar to that of control iPS cells (Figure 4D), suggesting that GPR161 may not play a critical role in iPS cell maintenance.

**Figure 4.**
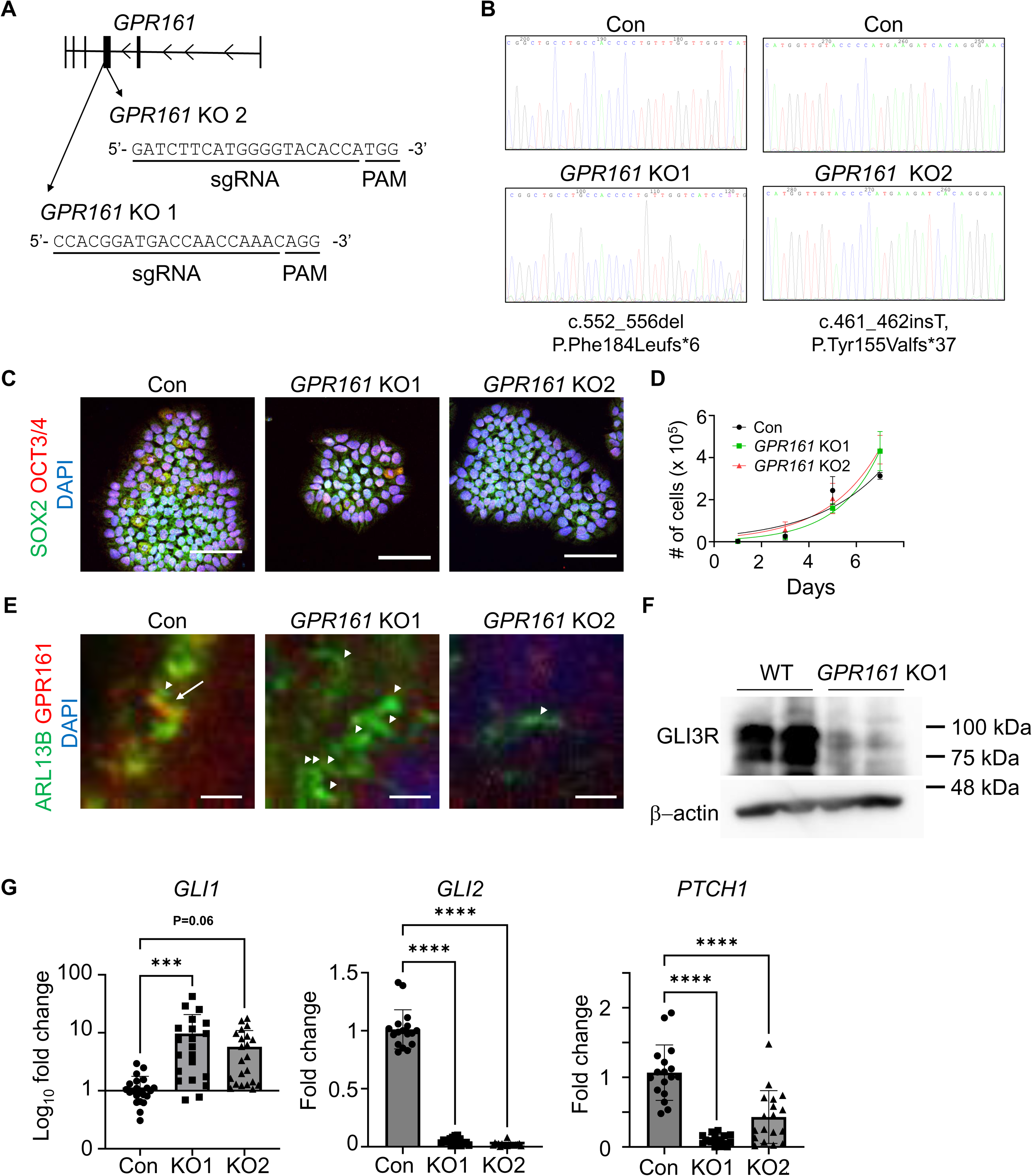
*GPR161* deletion induces disrupted SHH signaling in human brain organoids. (**A**) A scheme of *GPR161* gene with CRISPR/Cas9 target sequences. (**B**) Sanger sequences of the CRISPR/Cas9-targeted loci in control, *GPR161* KO1 and *GPR161* KO2 iPS cells (#51). (**C**) SOX2 and OCT3/4 expression in control, *GPR161* KO1 and *GPR161* KO2 iPS cells. (**D**) Cell growth curves of control, *GPR161* KO1 and *GPR161* KO2 iPS cells. N = 3 technical replicates. (**E**) GPR161 localization in ARL13B-positive primary cilia were shown in the ventricular zone of control, but not in *GPR161* KO1 and KO2 brain organoids. The arrow heads indicate primary cilia. The arrow indicates GPR161 localization in a primary cilium. (**F**) Western blot analysis revealed the reduction of GLI3 repressor (GLI3R) in *GPR161* KO1 brain organoids. β-actin is a loading control. (**G**) qRT-PCR revealed altered expression patterns of SHH target genes in *GPR161* KO1 and *GPR161* KO2 brain organoids during proliferation period. Each dot indicates mRNAs derived from 10–30 brain organoids. N = 17–22 accumulation of brain organoids. Scale bars are (**C**) 100 µm and (**E**) 1 µm.

To examine the function of GPR161-mediated signaling in neural stem cells, we generated dorsal cortical brain organoids using *GPR161* KO iPS cells using the guided dorsal cortical organoid differentiation protocol. We first confirmed that GPR161 was absent from primary cilia in the ventricles of brain organoids after 1 month of culture (Figure 4E). Consistent with previous studies, we showed that *GPR161* deletion decreased GLI3 repressor formation (Figure 4F). The deletion of *GPR161* induced the upregulation of *GLI1* but downregulation of *GLI2* and *PTCH1* in neural stem cells (Figure 4G). These data indicate that GPR161 regulates SHH signaling by inducing production of the GLI3 repressor, consistent with previous reports ^12,13,73^.

### Deletion of *GPR161* induces ventralization of brain organoids

Dominant negative variants of *GPR161* have been identified in spina bifida patients, indicating the critical role of GPR161 during brain development ^74^. We examined the role of GPR161 in the formation of dorsal/ventral neural stem cells in dorsal cortical brain organoids. Almost all cells in the ventricular zone expressed SOX2, indicating the cells in the ventricular zones are neural stem cells (Figure 5A, B). Although we employed a method to generate dorsal cortical brain organoids using dual SMAD inhibitors ^59^, *GPR161* KO iPS cells-derived brain organoids failed to generate PAX6-positive or PAX7-positive dorsal neural stem cells, indicating the failure in the generation of dorsal neural stem cells (Figure 5C-F). We observed OLIG2-expressing neural stem cells occasionally in both control and *GPR161* KO brain organoids (Figure 5G, H). Notably, we observed the increased number of NKX2.2-positive and FOXA2-positive ventral neural stem cells in *GPR161* KO brain organoids as compared to control brain organoids (Figure 5I–L). These data indicate that the deletion of *GPR161* prevents the formation of dorsal neural stem cells and induces ventral neural stem cell identity during cortical brain organoid formation using the guided differentiation protocol.

**Figure 5.**
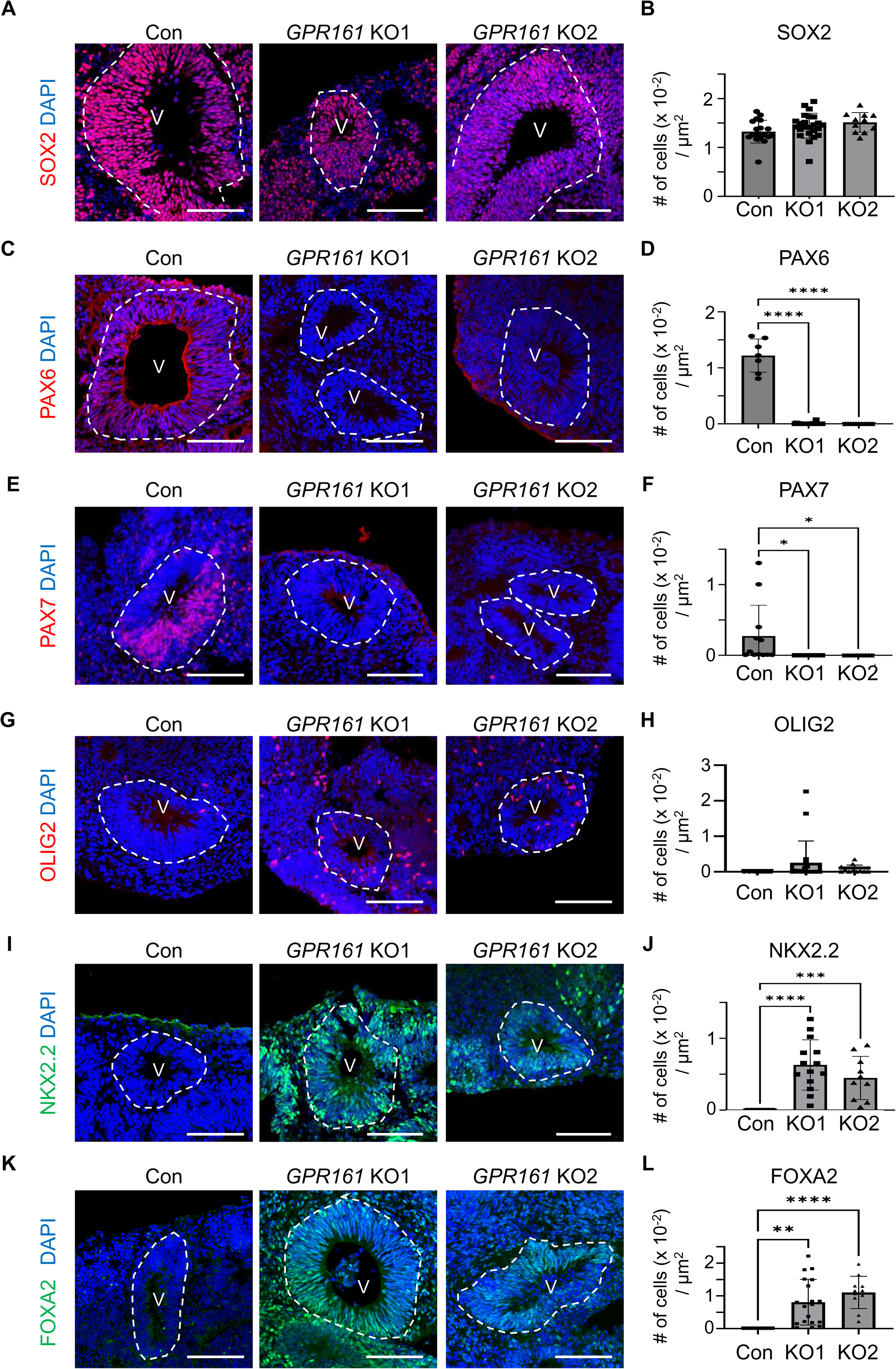
Deletion of *GPR161* induces ventralized brain organoids. (**A**, **B**) SOX2-positive pan neural stem cells were observed in control, *GPR161* KO1 and *GPR161* KO2 brain organoids. White dotted lines indicate ventricular zones. V, ventricle. N = 11–24 ventricular zones of brain organoids. (**C**, **D**) Reduced PAX6-positive dorsal neural stem cells in *GPR161* KO1 and *GPR161* KO2 brain organoids. N = 7–11 ventricular zones of brain organoids. (**E**, **F**) Reduced PAX7-positive dorsal neural stem cells in *GPR161* KO1 and *GPR161* KO2 brain organoids. N = 10–14 ventricular zones of brain organoids. (**G**, **H**) No difference in the number of OLIG2-positive neural stem cells in control, *GPR161* KO1 and *GPR161* KO2 brain organoids. N = 11–19 ventricular zones of brain organoids. (**I**, **J**) Increased NKX2.2-positive ventral neural stem cells in *GPR161* KO1 and *GPR161* KO2 brain organoids. N = 10–14 ventricular zones of brain organoids. (**K**, **L**) Increased FOXA2-positive ventral neural stem cells in *GPR161* KO1 and *GPR161* KO2 brain organoids. N = 11–19 ventricular zones of brain organoids. * indicates p < 0.05, ** indicates p < 0.01, *** indicates p < 0.001 and **** indicates p < 0.0001. Scale bars are (**A**, **C**, **E**, **G**, **I** and **K**) 100 µm.

### cAMP-mediated signaling rescues dorsalization of *GPR161* KO neural stem cells

Previous studies showed that GPR161 utilized the GaS-cAMP-PKA-GLI3 axis to regulate the SHH signaling pathway ^13^. To examine whether cAMP-mediated signaling is critical for dorsal/ventral determination, we treated *GPR161* KO brain organoids with forskolin, a chemical that increases cAMP levels through adenylyl cyclase activation. We examined the effect of forskolin at various concentrations and periods, and found that the treatment with 500 nM forskolin for the entire period most effectively promoted dorsalization of neural stem cells in *GPR161* KO brain organoids (Supplemental Figure 3A, B). The addition of forskolin did not affect the numbers of PAX6-dorsal neural stem cells and NKX2.2-or FOXA2-ventral neural stem cells in control brain organoid (Figure 6A-D). Notably, forskolin treatment significantly rescued the number of PAX6-dorsal neural stem cells, and depleted the numbers of NKX2.2-or FOXA2-ventral neural stem cells in *GPR161* KO brain organoids (Figure 6A-D). Forskolin treatment partially altered SHH target gene expression patterns (Supplemental Figure 3C-E). Importantly, the amount of GLI3 repressor was rescued by forskolin treatment in *GPR161* KO brain organoid, indicating the critical role of cAMP in GLI3 repressor formation (Figure 6E). The re-analysis of previously published CUT&Tag data using cerebral organoids ^75^ showed that GLI3 directly interacts with the *NKX2.2* locus, indicating that GLI3 directly inhibits *NKX2.2* (Figure 6F). These data indicate that the GPR161-cAMP axis is critical for dorsal/ventral fate choice of neural stem cells during brain organoid formation.

**Figure 6.**
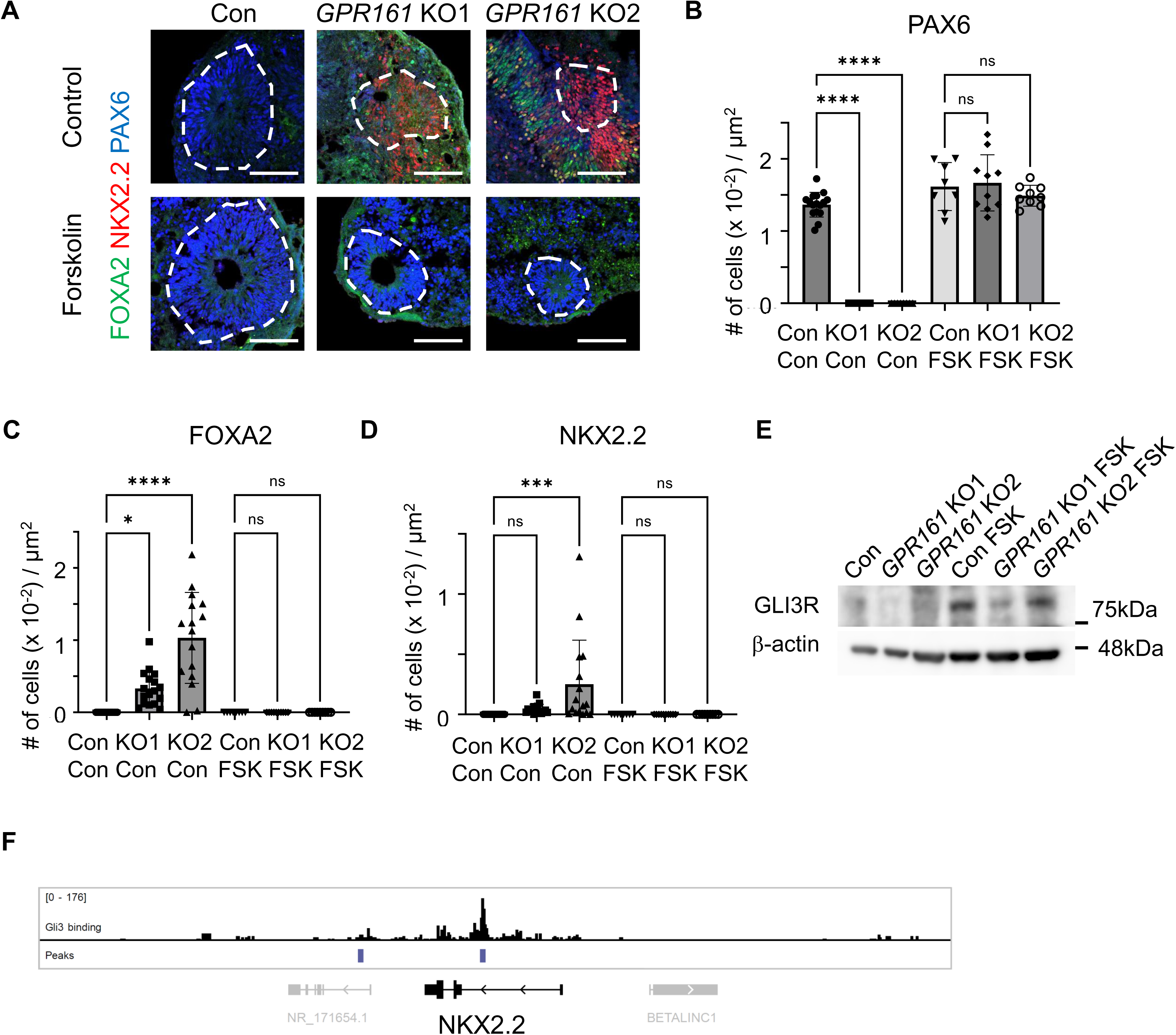
Forskolin treatment rescues dorsalization of *GPR161* KO brain organoids. (**A**-**D**) Forskolin treatment rescued dorsalization of neural stem cells in *GPR161* KO brain organoids. N = 5–17 ventricular zones of brain organoids. (**E**) Western blot analysis revealed that the reduction of GLI3 repressor (GLI3R) form in *GPR161* KO brain organoids were rescued by forskolin treatment. β-actin is a loading control. (**F**) CUT&Tag data of GLI3 binding around the *NKX2.2* gene locus in 23-day-old human cerebral organoids ^75^. Signal tracks (upper, black), called peaks (middle, blue), and genes (lower, gray and black) are shown. * indicates p < 0.05 and **** indicates p < 0.0001. Scale bars are (**A**):100 µm.

### Ciliary cAMP-mediated signaling rescues dorsalization of *GPR161* KO neural stem cells

One of the functions of primary cilia is the regulation of cAMP levels in the cilia for the SHH signaling pathway ^22,76,77^. To examine whether an increase in ciliary cAMP is sufficient to determine the dorsal/ventral fate of neural stem cells in *GPR161* KO brain organoids, we generated cells capable of modulating ciliary cAMP levels in the *GPR161* KO condition by inserting the *CAG-ARL13B-bPAC-EGFP* construct into the *AAVS1* locus on chromosome 19 of *GPR161* KO iPS cells (Figure 7A, Supplemental Figure 4A; hereinafter referred to as *ciliary-bPAC-GPR161* KO). bPAC is a protein that can produce cAMP when exposed to light at the 470 nm wavelength, and *ciliary-bPAC-EGFP* is a fusion protein that allows bPAC to localize to the cilia ^22^. We then generated the brain organoids and confirmed that bPAC-EGFP was enriched in ARL13B-positive primary cilia of neural stem cells in the ventricular zones of brain organoids (Figure 7B). To examine whether ciliary-cAMP-mediated signaling is critical for dorsal/ventral determination, *ciliary-bPAC-GPR161* KO brain organoids were exposed to programmed light stimulation for two weeks (Supplemental Figure 4B–D). Dark or light exposure conditions did not affect the dorsal/ventral fate of control and *GPR161* KO brain organoids (Figure 7C, D). *Ciliary-bPAC-GPR161* KO brain organoids in dark condition contained both PAX6-and NKX2.2-positive neural stem cells, indicating a low-level bPAC activity in the brain organoids in dark (Figure 7C, D). Notably, light exposure reduced the number of NKX2.2-positive neural stem cells in *ciliary-bPAC-GPR161* KO brain organoids (Figure 7C, D). These data indicate that an increase in ciliary cAMP levels following light exposure was sufficient to rescue dorsalization of *GPR161* KO brain organoids (Figure 7E).

**Figure 7.**
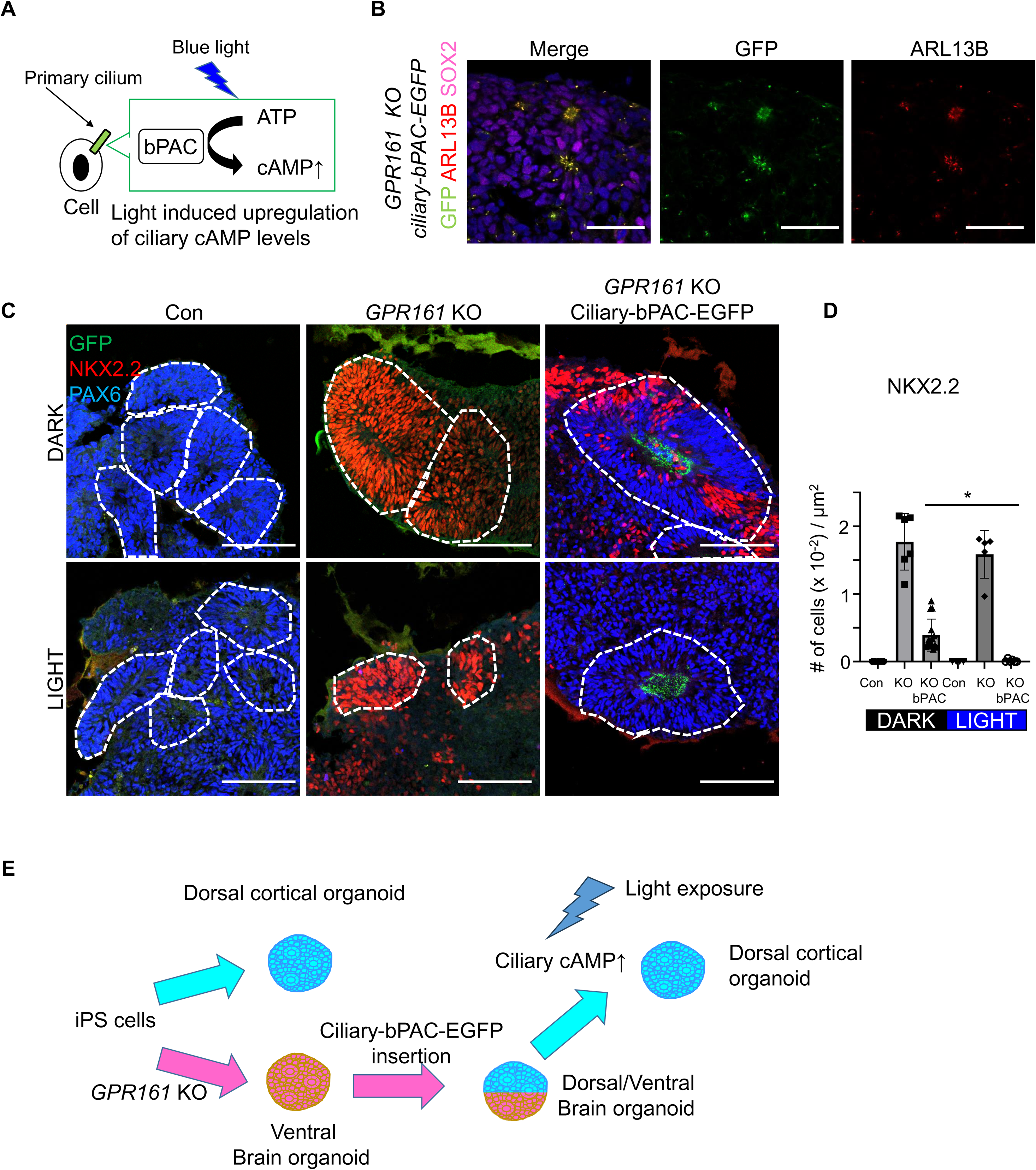
The elevation of ciliary cAMP levels rescues dorsalization of *GPR161* KO brain organoids. (**A**) A scheme of optogenetics assay in ciliary-bPAC-EGFP-expressing iPS cells. (**B**) Primary cilia in the ventricular zone of the brain organoids derived from ciliary-bPAC-*EGFP* expressing *GPR161* KO iPS cells. Ciliary-bPAC-EGFP was observed only in the ARL13B-positive primary cilia. (**C, D**) The elevation of ciliary cAMP levels rescued dorsalization of ciliary-bPAC-*EGFP* expressing *GPR161* KO brain organoids following light exposure. White dotted lines indicate ventricular zones. N = 4–15 ventricular zones of brain organoids. (E) Summary of the phenotypes of brain organoids in optogenetics assay. * indicates p < 0.05. Scale bars are (**B** and **C**) 100 µm.

## Discussion

Previous studies have shown that primary cilia are extended from neural stem cells in the mouse cortex and brain organoid models, but little is known about the mechanisms by which primary cilia play role in the fate decision of neural stem cells in human. Here, we have uncovered that the deficiency of ARL13B leads to partial ventral differentiation of neural stem cells in dorsal cortical organoids despite using the guided differentiation protocol (Figure 2 and 3). Additionally, our study identified the induction of ventralization of neural stem cells in *GPR161* deficient dorsal cortical organoids using the guided differentiation protocol (Figure 4 and 5). Mechanistically, we revealed that both ARL13B and GPR161 regulate the SHH signaling pathway, partly through GLI3 repressor formation (Figure 3 and 4). Remarkably, the upregulation of cAMP by forskolin treatment and ciliary cAMP activation by optogenetics successfully rescued the dorsalization of *GPR161* KO neural stem cells. The rescue was partly mediated by GLI3 repressor formation and the direct negative regulation of NKX2.2 (Figure 6 and 7). We and others previously showed the role of *Gpr161* in neural tube formation and cortical development in mice ^12,13,78,79^. Our current study confirmed that mouse and human share the same mechanisms for dorsal/ventral fate decision during brain development. It is important to note that an increase in ciliary cAMP levels alone was sufficient to rescue the dorsalization in *GPR161* KO brain organoids (Figure 7). In agreement with our study, inhibiting adenylyl cyclase maturation in mice disrupts ciliary cAMP signaling and causes neural tube ventralization ^21,80^. These compelling data suggest that the ciliary cAMP signaling pathway holds promise as a potential drug target for addressing neural tube closure and microcephaly.

### An enhanced protocol to generate dorsal cortical organoid

SHH signaling has critical roles in the cortical development. Indeed, previous studies have showed that the activation of SHH signaling induces ventralization of brain organoids ^38,57^. This hampers the analysis of SHH signaling in the dorsal cortical development using human brain organoids, since high SHH signaling prevents dorsal cortical organoid formation. To overcome the issue, we treated ventralized brain organoids with forskolin to change the dorsal/ventral fate of neural stem cells. Forskolin treatment promotes an increase in cAMP levels and induced dorsal brain organoids in the *GPR161*-deficient condition. Mechanistically, forskolin upregulates cAMP, which affects GLI3 cleavage to form GLI3 repressor. GLI3 directly binds to NKX2.2, indicating that GLI3 repressor formation is critical for NKX2.2-ventral neural stem cells.

### The ARL13B-GPR161-GLI signaling axis regulates dorsal/ventral fate of neural stem cells

SHH signaling determines dorsal/ventral fate of neural tube in various animal models ^41,81,82^. Our data suggest that both ARL13B and GPR161 contribute to the regulation of the SHH signaling pathway. Deletion of *ARL13B* induced partial ventralization, whereas deletion of *GPR161* induces strong ventralization. The upregulation of a SHH target gene GLI1 in both *ARL13B* and *GPR161* KO brain organoids suggests that SHH signaling is one of the signaling pathways to determine ventralization. However, it remains unclear how *ARL13B* deletion results in milder ventralization of neural stem cells compared to *GPR161* deletion. One possible reason is that ARL13B affects more than just the SHH signaling. It is well-known that primary cilia serve as a signaling hub for various signaling pathways, not only SHH signaling ^20,21,83–85^. Currently, the specific roles of ARL13B in primary cilia in human neural stem cells is uncertain, and unveiling these mechanisms will be interesting to reveal other functions of primary cilia in human brain development.

### ARL13B regulates multiple aspects of SHH signaling

ARL13B is a small GTPase with multiple function in primary cilia. Our data revealed that ARL13B is a critical factor to determine the morphology of primary cilia and SHH signaling in the dorsal cortical organoids (Figure 2). Interestingly, deletion of *ARL13B* induces partial ventralization of neural stem cells (Figure 3). Previous mouse studies showed that the disruption of *ARL13B* induced ventralization of the neural tube with increased SHH target gene expression ^62,86–88^. Deletion of *Gli2* in *Arl13b* mutant mice showed the rescued phenotype of dorsalization of the neural tube and optic vesicle morphogenesis, indicating the importance of upregulated SHH signaling in *Arl13b*-deleted condition ^62,89^. These data indicate that increased SHH signaling determines the fate of ventralization of neural tube. However, interestingly, SHH-activated medulloblastoma in mice showed that the tumor formation and the activity of the SHH signaling pathway were decreased following *Arl13B* deletion ^61^. In addition, treatment of Shh or the smoothened agonists on *Arl13B*-deleted cultured cells showed disrupted upregulation of the SHH signaling pathway ^61,88,90^. These data indicate that ARL13B has double-edged functions for SHH regulation in physiological and pathological conditions. It is interesting to examine if ARL13B also has a double-edged function in human cells, such as human SHH subtype medulloblastoma.

### Ciliary signaling determines dorsal/ventral fate of neural stem cells in brain organoid models

The importance of primary cilia in human brain development is evident in heterogeneous human diseases called ciliopathies, including Joubert syndrome, Bardet-Biedl syndrome, neural tube defects, and multipole mental retardation ^20,25^. The roles of primary cilia in neural stem cells are well studied in mice. However, it has been difficult to examine the role of ciliary signaling in human brain until the era of brain organoid research. ^24,91–93^. Recent studies showed that brain organoids contain primary cilia in the ventricular zone ^15^. In agreement with our study, other ciliary genes such as *INPP5E* and *GLI3* are involved in dorsal/ventral determination of neural stem cells in human brain organoids, using a non-guided differentiation protocol ^57,75^. The non-guided protocol allows to generate both dorsal/ventral fate of neural stem cells in a brain organoid, therefore the strength of SHH signaling was not clear. Another study also revealed the importance of primary cilia in the differentiation of neural stem cells into basal progenitor cells ^40^. These data indicate that primary cilia are a critical component of human neural stem cells and could be a future druggable target for brain developmental diseases.

### Limitation of the study

Our optogenetic model showed a certain amount of dorsalization of *GPR161* KO brain organoid without light exposure, suggesting some level of leaked cAMP at dark condition. Although our data clearly showed that light exposure eliminated NKX2.2-positive ventralized neural stem cells, it is important to generate light-sensitive optogenetic model with no leakage for further experiments.

## Materials and Methods

### iPS cell culture

Previously characterized human induced pluripotent stem cells (iPS cells; 201B7 (Riken) and #51 (also known as Windy; Dr. Umezawa)) were used in the current study ^94–97^. iPS cells were maintained in Stemfit (Ajinomoto, RCAK02N) media with iMatrix-511 (Takara Bio, T311). The media was changed every one to three days for routine maintenance. iMatrix-511 was used in an uncoated manner (0.2 ug/cm^2^), as previously described ^98^. For the routine maintenance, iPS cells were passaged every five to seven days using 0.5 mM EDTA and TypLE express enzyme (Gibco, 12604021). For passage, cells were rinsed once with 0.5 mM EDTA, and then incubated with TypLE express enzyme/0.5 mM EDTA (1:1 ratio) for five minutes at 37 C. Cells were then collected in a tube and centrifuged at 190 × *g* for three minutes. Cell numbers were counted and cells were seeded at a density of 750 cells/cm^2^ for passage. After passage, the culture media were supplemented with 10 µM Y-27632 (Chem scene, CS-0878) for the first 24 hours. 150 ul of Stem Cell Banker (Takara, CB045) was routinely used for cryopreservation of 1-50 × 10^4^ cells. Cells were used for experiments between passages 40-60. Cells were tested for mycoplasma by PCR. Primers are listed in Supplemental Table 1.

### Generation of *ARL13B* and *GPR161* knockout iPS cells

Clustered regularly interspaced short palindromic repeats (CRISPR)/Cas9-mediated genome editing was used to delete *ARL13B* or *GPR161* in iPS cells. For the preparation of ribonucleoprotein (RNP) complex, 2 µl of 100 µM CRISPR crRNA (Integrated DNA Technologies) and 2 µl of 100 µM Alt-R tracrRNA were mixed with Atto^TM^ 550 (Integrated DNA Technologies, 1075927), followed by incubation at 95°C for 5 minutes and then at room temperature to form a single guide RNA (sgRNA) complex. One µl of 62 µM Alt-R S.p. HiFi Cas9 protein (Integrated DNA Technologies, 1081060) was added to the sgRNA complex and incubated in the dark at room temperature for 30 minutes to form the RNP complex. Subsequently, 0.5–1L×L10^6^ iPS cells were harvested and electroporated with 5 µl of the prepared RNP complex dissolved in 100 µl electroporation solution (6.6 mM ATP, 11.8 mM MgCl_2_-6H_2_O, 147 mM KH2PO_4_, 23.8 mM NaHCO_3_, 3.7 mM glucose, pH7.4) using Nucleofector 2b (Program #B-016; Lonza) ^99^. Cells were centrifuged in DMEM/F12 (Invitrogen, 11330-032) and plated for cell culture. The next day, cells were harvested and stained for DAPI (Sigma-Aldrich, D8417) in Stemfit media. DAPI-negative and Atto^TM^ 550-positive iPS cells were analyzed or isolated using a cell sorter (FACSCanto II, BD Biosciences; FACSAria II; BD Biosciences, USA; FACSAria III; BD Biosciences, USA) and plated in a dish with the cell density of 100-200 cells per 10 cm^2^ dish. Following day, the cells were observed and confirmed iPS cells were clonally cultured. Non-clonal colonies were manually scratched with a 20 µl tip and removed from the plate under a microscope (Olympus, CKX53). Five to Seven days after sorting, cells were treated with 0.5 mM EDTA for three minutes and three ml of DMEM/F12 was added to the plate. Single clones were picked manually with a 20 µl tip and plated into a 24-well plate, and cultured for 7-14 days. After the expansion of the clones, cells were harvested. Half of the cells were stored for DNA extraction while the other half was frozen for passage. Genomic DNA was extracted in Proteinase K lysis buffer (10 mM Tris-HCl pH8.8, 50 mM KCl, 2 mM MgCl_2_, 0.45% NP-40, 0.45% Tween 20, 0.2 mg/ml Proteinase K), and PCR was performed to examine the sequence of the target region to obtain knockout cell lines. The cell lines generated in this study are available from corresponding authors upon reasonable request, after obtaining permission from Riken and of the National Center for Child Health and Development. Primer sequences are listed in Supplemental Table 1.

### Construction of *pAAVS1-CAG-ARL13B-bPAC* plasmid

*pAAVS1-CAG-ARL13B-bPAC* plasmid was generated from various plasmids. First, *pAAVS1-CRYS-ARL13B-bPAC-EGFP* plasmid was constructed using In-Fusion HD Cloning Kit (Takara, 639650). The backbone sequence was cloned from *pAAVS1-P-CAG-mCh* plasmid (Addgene 80492^100^) using the primers (YH146 and YH 147, see Supplemental Table 1) and Q5 High-Fidelity DNA Polymerase (NEB, M0491L). The insertion sequences were cloned from *pgLAP5-CRYS-ARL13B-bPAC-EGFP* plasmid (gift from Dr. Jeremy Reiter at UCSF ^22^) using the primers (YH144 and YH145). *pAAVS1-CMV-ARL13B-bPAC-EGFP* plasmid was constructed using In-Fusion HD Cloning Kit. The backbone sequence was cloned from *pAAVS1-CRYS-ARL13B-bPAC-EGFP* plasmid using the primers (YH162 and YH163). CMV promoter was cloned from pLJM1-Empty plasmid (Addgene 91980^101^) using the primers (YH160 and YH161). *pAAVS1-CAG-ARL13B-bPAC-EGFP* plasmid was constructed using In-Fusion HD Cloning Kit. The backbone sequence was cloned from *pAAVS1-CAG-EGFP* (Addgene 80491; ^100^) plasmid using the primers (pAAVS1-after chimeric intron-F and pAAVS1-prior to EGFP-R). The insert (ARL13B-bPAC-EGFP) sequence was cloned from *pAAVS1-CMV-ARL13B-bPAC-EGFP* plasmid using the primers (chickenintron-ARL13B-F and linker-R).

### Generation of *ciliary-bPAC-GPR161* KO iPS cells

To generate *ciliary-bPAC*-*GPR161* KO iPS cells, we utilized a published protocol to insert one copy of the gene of interest into the *AAVS1* locus on the chromosome 19 ^100^. *GPR161* KO iPS cells were plated onto 24-well plates in 6,000 cells/well density. The next day, cells were transfected with pXAT2 (Addgene #80494^100^) and *pAAVS1-CAG-ARL13B-bPAC-EGFP* plasmids using Lipofectamine Stem Transfection Reagent (Thermofisher, STEM00001). The cells were cultured until confluent and EGFP-positive cells were sorted using FACS. The cells were plated onto 60 cm dish, and then cultured until confluency and then EGFP-positive cells were sorted using FACS for the second time. The cells were plated onto 60cm dish and then used for experiments or cryopreserved.

### Generation of brain organoids from iPS cells

To generate brain organoids, we used a dual SMAD inhibitor protocol with slight modifications ^59,102^. We seeded 3,000 iPS cells into ultra-low attachment-coated 96-well U-bottom plates (Corning, 7007; ThermoFisher Scientific, 174925) and cultured them in the induction medium (DMEM/F12 containing 20% knockout serum replacement (ThermoFisher Scientific, 10828010), 1 mM MEM non-essential amino acids (Gibco, 11140), 2 mM L-glutamine (Nacalai, 16948-04), 100 µM 2-mercaptoethanol (Wako, 133-14571), 100 nM LDN-193189 (Cayman Chemical, 19396), 10 µM SB-431542 (Cayman Chemical, 13031), and 2 µM XAV939 (Cayman Chemical, 13596)) with 50 µM Y-27632 (for the first 24 hours) for the first six days with replacement every two-three days. To obtain a large number of organoids for protein/RNA collection, 188,000 iPS cells were seeded into EZ-sphere 24-well plates (approximately 400 cells per microwell; Iwaki, 4820-900SP). At day 6, the induction medium was removed and 2% of Matrigel (Corning, 354234) was added to the bottom of the plate and cultured in the organoid culture medium 1 (50% DMEM/F12, 50% MACS Neuro Medium (Miltenyi Biotec, 130-093-570), 1 mM MEM non-essential amino acids, 2 mM L-glutamine, 50 µM 2-mercaptoethanol, 400 nM insulin (Nacalai, 12878-86), 0.5× N-2 MAX Media Supplement (R and D systems, AR009), 100 U/ml penicillin/streptomycin (Nacalai, 26253-84) and 0.5× MACS NeuroBrew21 supplement without vitamin A (Miltenyi Biotec, 130-097-263)) for 8 days replaced the media every two to three days. Organoids were transferred to six-well plates on day 14. Six-well plates can be pre-coated with 2-hydroxyethyl methacrylate solution (1.2 g of 2-hydroxyethyl methacrylate (Sigma-Aldrich, P3932) in 50 ml of 95% ethanol) for low binding condition (optional). To mature the organoids, the spheroids were cultured in organoid culture medium 2 (50% DMEM/F12, 50% MACS Neuro Medium, 1 mM MEM non-essential amino acids, 2 mM L-glutamine, 50 µM 2-mercaptoethanol, 400 nM insulin, 0.5× N2 supplement, 100 U/ml penicillin/streptomycin, 0.5× MACS NeuroBrew21 with vitamin A (Miltenyi Biotec, 130-093-566), and 200 µM L-ascorbic acid (Sigma-Aldrich, A92902)) from day 14 up to one month. The plates were rotated on an orbital shaker (TAITEC, CS-LR) at 70 rpm. The culture media were replaced every one to four days.

### Generation of neural stem cells for 2D culture

To generate iPS cells-derived neural stem cells was performed as previously described ^97,103,104^. 188,000 iPS cells were seeded into EZ-sphere 24-well plates and cultured in the induction media with 50 µM Y-27632 (for the first 24 hours) for the first six days with replacement every two-three days (approximately 400 cells per microwell). On the sixth day, spheres were seeded into six-well plates that had been coated with 20 ng/ml laminin (Corning, 354232) in Poly-D-Lysine (R and D systems, 343-100-01) for 1 hour at 37°C. The attached spheres were cultured in the Neural stem cell media (50% DMEM/F12 and 50% MACS Neuro Medium containing 1 mM MEM non-essential amino acids, 2 mM L-glutamine, 0.5× N-2 MAX media supplement, 0.5× MACS NeuroBrew21, 20 ng/ml FGF-2 (Gift from Dr. Spees), 20 ng/ml EGF (Almone Labs, E-100), 100 U/ml penicillin, and 100 µg/ml streptomycin). The sphere-derived monolayer cells were cultured until the neural rosette formation, with the replacement of media every two-three days. Subsequently, cells were dissociated, passaged into a new Poly-D-Lysin/laminin-coated dish with Y-27632, and/or cryopreserved in Stem Cell Banker.

### Sample processing, antibodies, immunostaining and microscopy

Brain organoids were fixed with 4% paraformaldehyde (Wako, 162-16065) in PBS for 3–4 hours at 4°C, and incubated in 30% sucrose (Nacalai, 09589-05) in PBS for 1 day at 4°C. The length of paraformaldehyde fixation is important for GPR161 immunohistochemistry. Brain organoids were mounted with OCT compound (Sakura Fintek Japan, 4583), and cut into 16 µm frozen sections. For immunostaining using frozen sections, the sections were incubated in PBS for 15 min to dissolve away the OCT. Sections were then blocked in blocking buffer (3% normal serum block (Biolegend, 927503), and 0.4% Triton X-100 (Sigma-Aldrich, 30-5140-5) in PBS) for 1 hour at room temperature. For OLIG2 staining, FBS blocking buffer was used (3% fetal bovine serum (FBS; Sigma-Aldrich, 172012), 0.4% Triton X-100 in PBS). Sections were incubated with primary antibodies against the following antigens overnight at room temperature: acetylated alpha-tubulin (Sigma-Aldrich,T6793), ARL13B (Proteintech, 17711-1-AP), ARL13B (Biolegend, 857602), β-actin (Santa Cruz, sc-69879), beta-catenin (Sigma-Aldrich, C2206), beta-tubulin III (Santa Cruz, sc-80005), BLBP (Millipore, ABN14), FOXA2 (HNF-3b; Santa Cruz, sc-101060), gamma-tubulin (Santa Cruz, sc-7787), gamma-tubulin (Santa Cruz, sc-51715), GLI3 (R and D Systems, AF3690), GPR161 (Proteintech, 13398-1-AP), NKX2.2 (Hybridoma Bank, 74.5A5-s), OCT3/4 (Santa Cruz, sc-5279), OLIG2 (R and D Systems, AF2418), PAX6 (BioLegend, 901301), PAX7 (Hybridoma Bank, Pax7), pH3 (Santa Cruz, sc-374669), Poly-E (Adipogen, GT335), and SOX2 (Santa Cruz, sc-365823). After three PBS washes, the sections were incubated in secondary antibodies (Alexa Fluor 488-, 555-, 594-, 647-conjugated secondary antibodies, Invitrogen) for 1 hour at room temperature. Cell nuclei were stained with DAPI. Slides were mounted with Fluoromount-G (Southern Biotech, 0100-01) or CC/Mount (Diagnostic Biosystems, K002) and images were acquired with following microscopes (Olympus FV3000, Olympus SpinSR, Nikon AXR, or Nikon A1).

### Immunocytochemistry of iPS cells

iPS cells were plated on a plastic cover glass (Sumitomo Bakelite, MS-92132Z) in a 24-well plate. To examine primary cilia, cells were starved for 24 hours in DMEM/F12. Cells were fixed with 4% paraformaldehyde for 15 minutes at room temperature. Cells were blocked in blocking buffer and stained for antibodies. The plastic cover glass was mounted on a slide and images were acquired with microscopes.

### Immunocytochemistry of neural stem cells

Neural stem cells were plated on a Poly-D-lysin/laminin coated-cover glass (Marienfeld Superior™ Borosilicate Glass Cover Glasses, Round, Paul Marienfeld GmbH, 0111520) in a 24-well plate. Cells were fixed with ice-cold Methanol for 15 minutes at 4°C. Cells were blocked in blocking buffer and stained for antibodies. The glass cover glass was mounted on a slide and images were acquired with microscopes.

### Cell Growth Curve

iPS cells were seeded in 24-well plates at a density of 2,500 cells/cm^2^. To quantify the number of cells, cells were treated with 200 µl of TrypLE/0.5 mM EDTA solution and pipetted. Ten µl of the solution was taken and the number of cells was quantified on a hematocytometer.

### RNA isolation and qRT-PCR

Total RNA was isolated from brain organoids using Isogene II (NipponGene, 311-07361) or RNA premium kit (Nippon Genetics, FG-81050), according to the manufacturer’s instructions. cDNA was generated using ReverTra Ace qPCR RT Master Mix with gDNA remover (Toyobo, FSQ-301), according to the manufacturer’s instructions. qRT-PCR was performed with Faststart universal SYBR green master (Sigma-Aldrich, 4913914001). Reactions were run using a real-time PCR machine (ThermoFisher, Quant Studio 12K Flex). Primer sequences are listed in the Supplemental Table 1.

### Western blot

To obtain protein extracts for western blot analysis, 10–30 brain organoids were rinsed twice with ice-cold PBS and frozen until use. The frozen samples were lysed with RIPA lysis buffer (150 mM NaCl (Sigma-Aldrich, 3014), 1% NP-40, 0.5% sodium deoxycholate (Sigma-Aldrich, D6750), 0.1% SDS (Sigma-Aldrich, 28-3260-5), 50 mM Tris-HCl pH7.4) containing 1% protease and phosphatase inhibitor cocktail (ThermoFisher, 78445) for 3–4 hours at 4°C. The lysate was centrifuged (15,000 × *g*, 4°C) for 30 min, and the supernatant was used for further analysis. 4× SDS sample buffer (0.225 M Tris-HCl pH6.8, 50% glycerol (Sigma-Aldrich, 12-1120-3), 5% SDS, 0.05% bromophenol blue (Sigma-Aldrich, 03-4140-3), 0.25 M DTT (Nacalai, 14128-62)) was added and the samples were boiled at 100°C for 5 min. Samples were subjected to 9% homemade SDS-polyacrylamide gel electrophoresis with running buffer (25 mM Tris, 192 mM glycine, 0.1% SDS), followed by transfer to polyvinylidene difluoride membranes (Millipore, IPVH00010) with transfer buffer (50 mM Tris, 192 mM glycine, 20% methanol (Sigma-Aldrich, 19-2410-4)). Non-specific binding was blocked with 5% skim milk in 0.1% Triton X-100 in PBS (PBST) for 1 hour at room temperature, and proteins were probed with primary antibodies diluted in 0.1% PBST at 4°C overnight. After washing with 0.1% PBST, the horseradish peroxidase (HRP)-labeled secondary antibodies against rabbit, mouse or goat IgG were probed for 1 hour at room temperature. Luminata Forte Western HRP Substrate (Millipore, WBLUF500) was used to detect the signal. All images were acquired from an Amersham Imager 680 (GE Healthcare). Subsequently, the blot was treated with Western Blot Stripping Buffer (ThermoFisher Scientific, 21059) for 10–15 min at room temperature. The same blot was re-probed with anti-β-actin antibody as a loading control. Primary and secondary antibodies are listed in Supplemental Table 1.

### Transmission electron microscopy (TEM)

Brain organoids were fixed in 2% glutaraldehyde (Electron Microscopy Sciences, 111-30-8), in 0.1 M phosphate buffer (PB; pH 7.4) at 4°C overnight, and post-fixed in 2% osmium tetroxide (Sigma-Aldrich, 1.24505) in 0.1 M PB (pH 7.4) at 4°C for 45 min. The samples were then dehydrated with a graded ethanol series and embedded in Quetol 812 epoxy resin (Nissin, 340-H) at 60°C for 48 hour. Ultrathin sections 80-100 nm thick were obtained with an Ultracut-UCT (Leica), stained with uranyl acetate (Merck, 8473) for 15 min, and stained with modified Sato’s lead solution ^105^ for 5 min. Transmission electron microscopy observations were performed using an electron microscope (JEM-1400 Plus; JEOL, Japan).

### Scanning electron microscopy (SEM)

Samples were cut in half or more using 76 razor (Nisshin EM, 4761) under a microscope, and then washed in Organoid 2 media in a 6-well plate for 30 minutes to clean the debris inside of the ventricles. Samples were pre-fixed in a fixative containing 2% glutaraldehyde in 0.1 M PB (pH 7.4) for more than 24 hours. After fixation, the samples were post-fixed with 2% osmium tetroxide in 0.1 M PB for 1 hour. Samples were placed in 25% DMSO solution for 30 min, replaced in 50% DMSO solution, cooled in liquid nitrogen. The samples were placed on a metal plate cooled with liquid nitrogen and stored until the sample was sufficiently frozen. The samples were collected in a cooled 50% DMSO solution and returned to room temperature. These specimens were dehydrated in ethanol, followed by critical point drying (Leica, CPD300). Samples were air dried in a hood, mounted on SEM specimen stubs, coated with a layer of sublimated OsO4 using an osmium plasma coater (OPC80N; Filgen, Nagoya, Japan). Images were acquired using a Field-Emission Scanning Electron Microscope (S-4700; Hitachi, Tokyo, Japan) at 5kV.

### CUT&Tag data of GLI3 binding

CUT&Tag data of GLI3 binding in 23-day-old cerebral organoids from human iPS cell line 409B2 was re-analyzed from published data ^75^. Raw Fastq files (downloaded from E-MTAB-12006) were processed through fastp (v0.23.2) ^106^ and mapped with Bowtie2 (v2.2.5) ^107^ to the human genome hg38. The resulting sam file was converted to the bigWig file using SAMtools (v1.15.1) ^108^ and deepTools (v3.5.1) ^109^. Peak calling was performed using HOMER (v4.11) ^110^. Signal tracks of GLI3 binding were visualized using IGV (v2.17.4) ^111^.

### Optogenetics

We generated a custom LED device to automatically regulate 470 nm LED exposure. The device consisted of Arduino Uno SMD R3 (Arduino, A000073), Gravity relay module (DFRobot, DFR0643), and LED device (Optocode, LEDB-SBOXHP). We performed optogenetic experiments in 96 low binding U bottom plate and 6-well plate using the custom LED device. During handling the cells, the cell culture room was kept dark using a 660 nm LED stand light (Optocode, LED660-100STND). The 470 nm LED was controlled by an Arduino Uno microcontroller, which was programmed with a custom script written in the Arduino Integrated Development Environment (Version 2.3.1; Supplemental Figure 4C). We stimulated cells with 470 nm blue light for two weeks using a pulsed pattern (500 ms ON, 2 min OFF) (Supplemental Figure 4D).

### Cell Quantification, Statistics and Software usage

To quantify the number of cells, we took pictures of the regions of interest and analyzed them using Fiji. Data from 3–20 ventricular zones in 3–10 brain organoids were used for the analysis with Fiji. To quantify the length of cilia of neural stem cells, we took pictures of the regions of interest using a 40× lens with 2× zoom (FV3000, Olympus) and quantify the length using image J manually. No blinding was performed. Sample sizes were based on our experience with these assays. All data in the figures are expressed as mean ± SD. To assess the statistical significance of differences among treatments, we often performed unpaired, two-sided Student’s *t*-tests that assumed unequal variances in treatments, one-way ANOVA with Sidak’s multiple comparisons tests and one-way ANOVA with Turkey multiple comparisons tests. GraphPad Prism (GraphPad, La Jolla, CA) were used for statistical analysis. Values of p < 0.05 were considered significant. In organoid experiments for quantifying specific area or cell number, n represents the number of individual ventricular zones. Organoid experiments were performed on biological replicates. For each independent experiment, the samples were newly generated from distinct passages of each iPS cell. We have used Chat GPT, DEEPL and Grammarly to correct English grammar of the current manuscript.

---

**Supplemental Figure 1.**
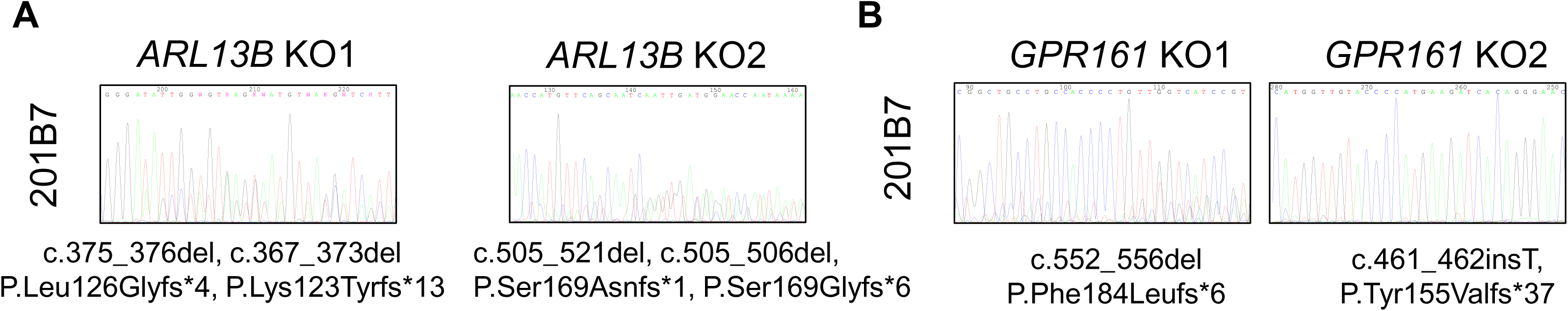
Sanger sequences of additional iPS cell lines. (**A**) Sanger sequences of the CRISPR/Cas9-targeted loci in control, *ARL13B* KO1 and *ARL13B* KO2 iPS cells (201B7 cells). (**B**) Sanger sequences of the CRISPR/Cas9-targeted loci in control, *GPR161* KO1 and *GPR161* KO2 iPS cells (201B7 cells).

**Supplemental Figure 2.**
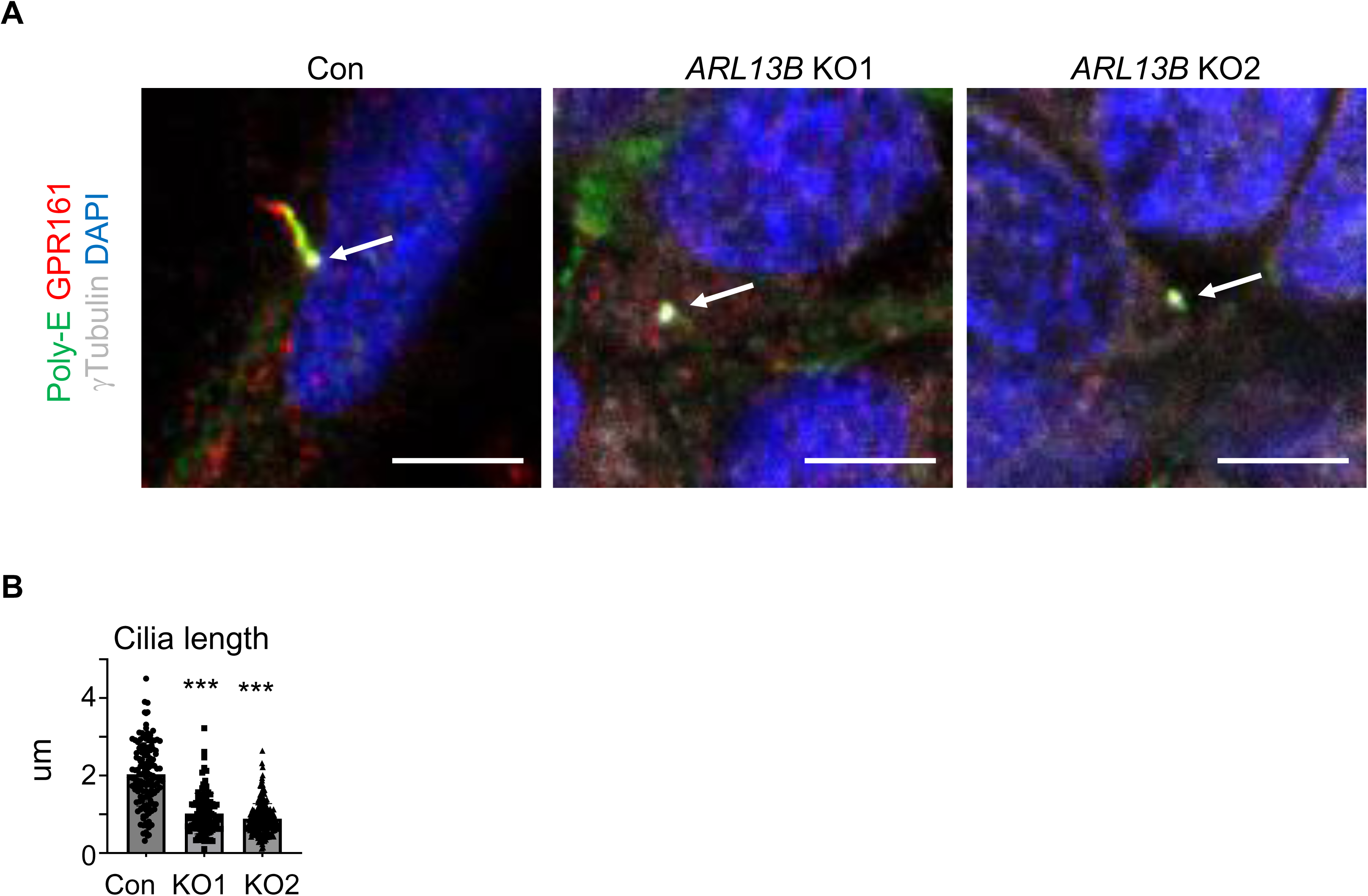
The ciliary length of neural stem cells. (**A**) Primary cilia were shown in control, *ARL13B* KO1 and *ARL13B* KO2 neural stem cells in 2D culture. Polyglutamylated tubulin (Poly-E) indicates primary cilia. Arrows indicate the basal bodies. Scale bars are 5 µm. (**B**) Comparison of the ciliary length in control, *ARL13B* KO1 and *ARL13B* KO2 neural stem cells in 2D culture. N = 108–196 cells in 2 independent experiments. *** indicates p < 0.0001.

**Supplemental Figure 3.**
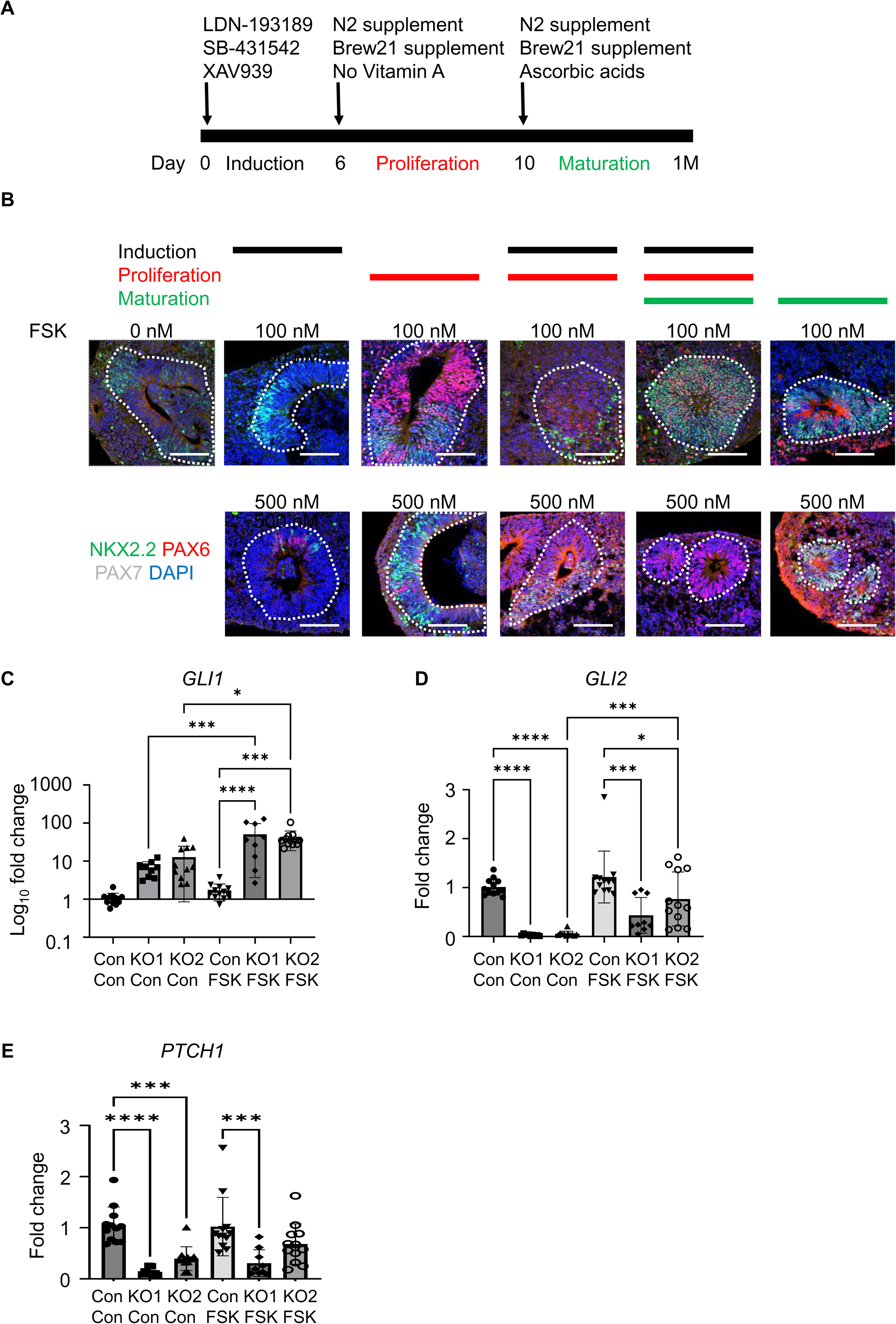
qRT-PCR of *GPR161* KO brain organoids following forskolin treatment. (**A**) A time schedule of brain organoid generation. (**B**) The induction of dorsalization of neural stem cells in *GPR161* KO1 brain organoids with various treatment schedules of forskolin (FSK). The black, red and green lines indicate the period with forskolin. White dotted lines indicate ventricular zones. **(C**–**E)** *GLI1*, *GLI2* and *PTCH1* expression levels in control, *GPR161* KO1 and *GPR161* KO2 brain organoids with/without forskolin (FSK) treatment. * indicates p < 0.05, ** indicates p < 0.01 and **** indicates p < 0.0001. scale bars are (**B**):100 µm.

**Supplemental Figure 4.**
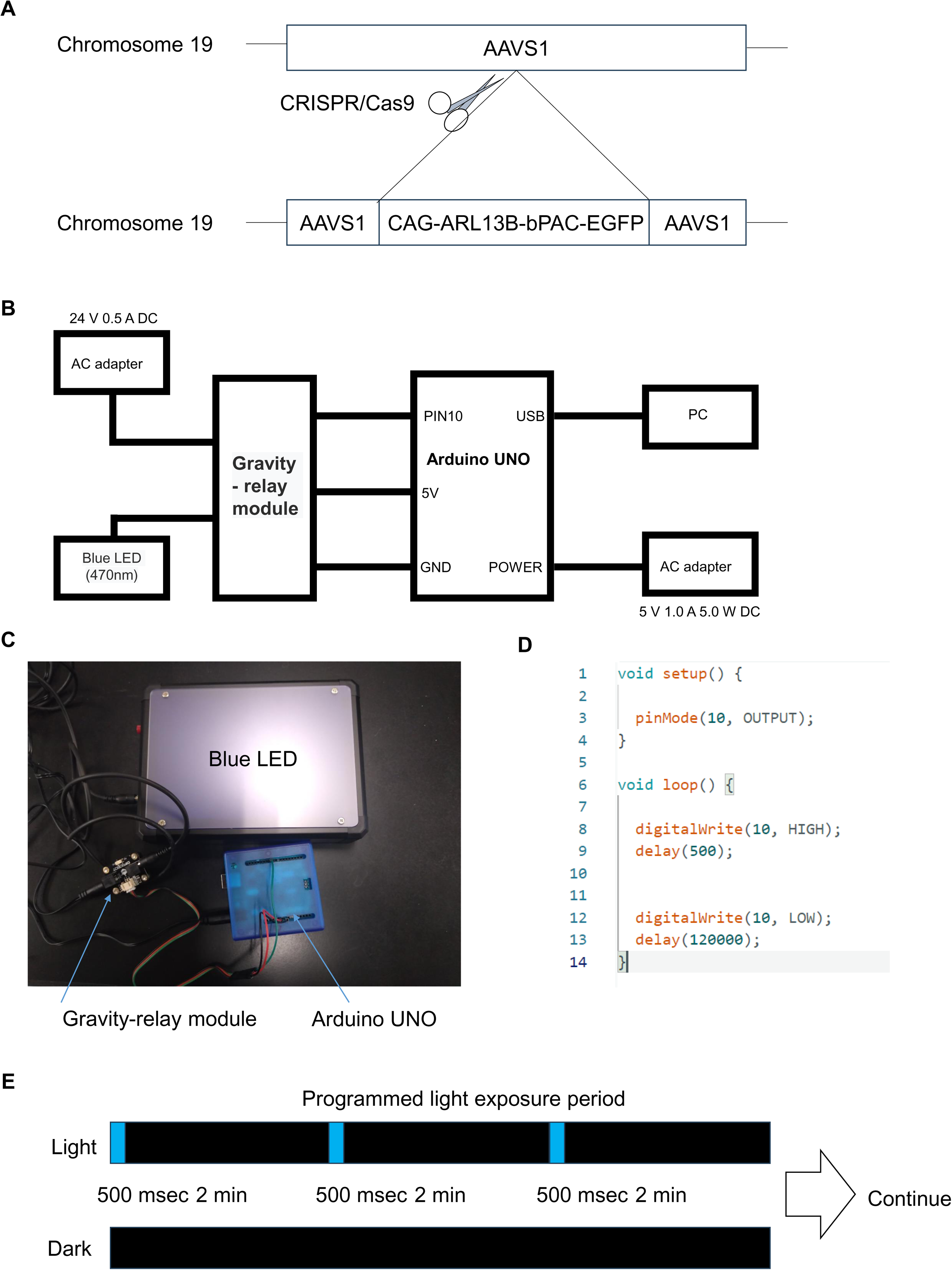
Optogenetic setup for the cell culture. (**A**) A scheme to generate gene knock-in at the safe harbor of chromosome 19. CAG-ARL13B-bPAC-EGFP was inserted into the *AAVS1* locus by CRISPR/Cas9. (**B**, **C**) The circuit and devices used in the current study. (**D**) Arduino programming for the current study. (**E**) A scheme of light exposure schedule.

**Supplemental Figure 5.**
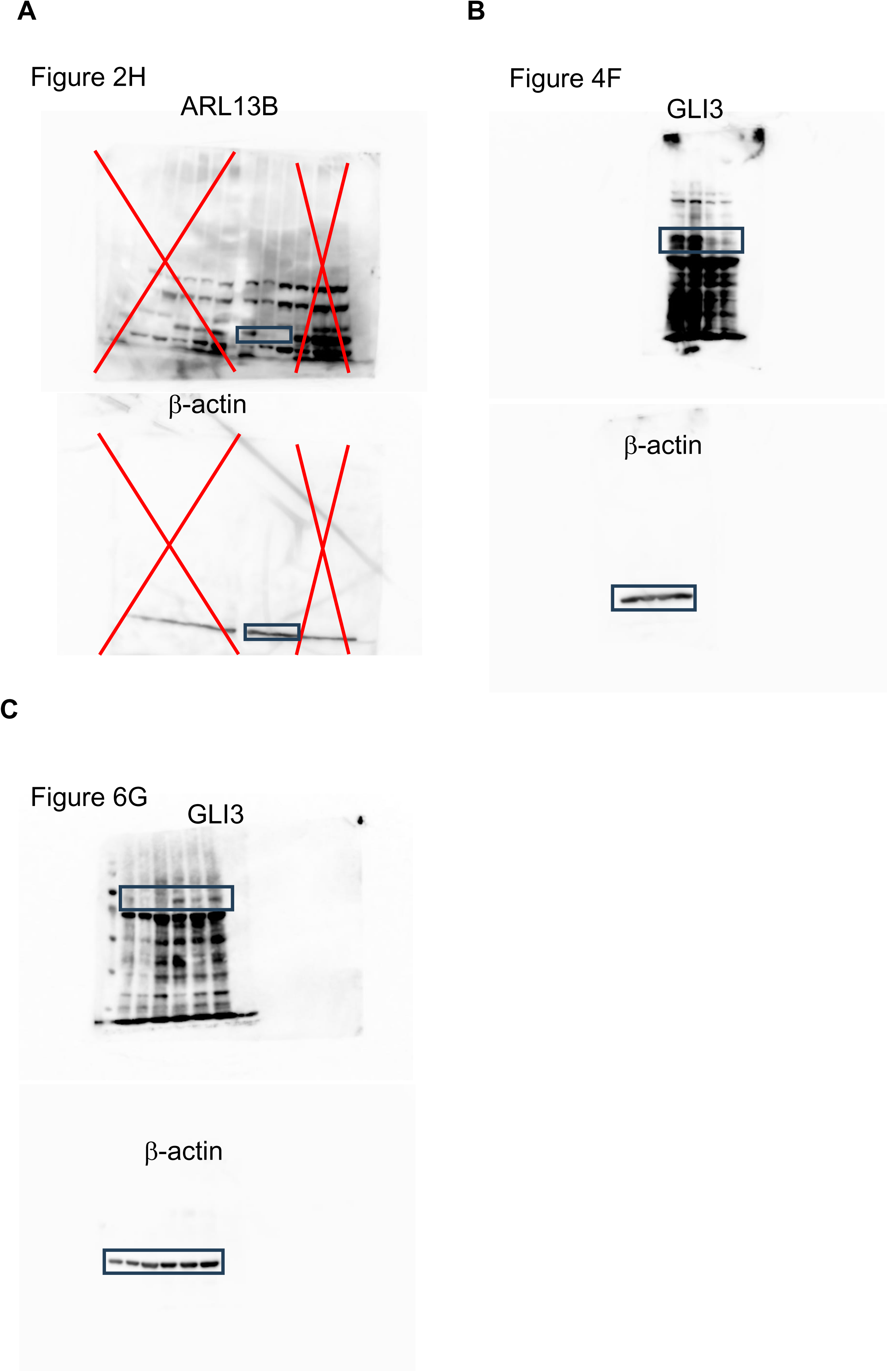
Non-cropped blots.

## Disclaimer

ISS and YK submitted a patent application about the dorsal brain organoid formation.

## Acknowledgments

Human iPS cells (Windy) were kindly provided by Dr. Umezawa of the National Center for Child Health and Development (Tokyo, Japan) and by Dr. Matsunaga of the Nagoya City University. Anti-NKX2.2 and anti-PAX7 antibodies were obtained from University of Iowa Hybridoma bank. We thank Dr. Jeffrey Spees at the University of Vermont for his kind gift of bFGF. We thank Dr. Jeremy Reiter at the University of California at San Francisco for his kind gift of bPAC vectors. We appreciate Drs. Sawamoto, Sawada and Saitoh of Nagoya City University for generously allowing us to use the electroporator. We thank Kohki Shimada of Shioji Junior High School for generation of the circuit and devices used for the optogenetic study. This work in supported by Daiko Foundation, The Hori Science and Arts Foundation, Mochida Foundation for Medical and Pharmaceutical Research, Takeda Science Foundation, iPS Academia Japan Inc., Leave a Nest Co. Ltd. (Ikeda Rika award), Grand-in-Aid for research in Nagoya City University (2013009), and Japan Society for the Promotion of Science Grant-in-Aid for Scientific Research (21K07803, 24K02425) to I.S.S., and Takeda Science Foundation and Japan Society for the Promotion of Science Grant-in-Aid for Challenging Research (20K21584) to Y.K. We thank to the technical assistance of the Research Equipment Sharing Center/Core Laboratory at Nagoya City University.

## AUTHOR CONTRIBUTIONS

I.S.S. and Y. K conceived the project, designed the experiments, analyzed most of the data, and wrote the paper. I.S.S. performed most of the experiments. A. G. analyzed data in Figure 4 and 5. M. I. analyzed data in Figure 1. H. T. performed scanning electron microscope and transmission electron microscope analysis. Y.H analyzed gene expression data.

## References

1. Rayon, T., Maizels, R. J., Barrington, C. & Briscoe, J. Single-cell transcriptome profiling of the human developing spinal cord reveals a conserved genetic programme with human-specific features. Development 148, (2021).

2. Zeng, B. et al. The single-cell and spatial transcriptional landscape of human gastrulation and early brain development. Cell Stem Cell 30, 851–866.e7 (2023).

3. Benito-Kwiecinski, S. et al. An early cell shape transition drives evolutionary expansion of the human forebrain. Cell 184, 2084–2102.e19 (2021).

4. Pinson, A. et al. Human TKTL1 implies greater neurogenesis in frontal neocortex of modern humans than Neanderthals. Science 377, (2022).

5. Casingal, C. R., Descant, K. D. & Anton, E. S. Coordinating cerebral cortical construction and connectivity: Unifying influence of radial progenitors. Neuron 110, 1100–1115 (2022).

6. Allen, D. E. et al. Fate mapping of neural stem cell niches reveals distinct origins of human cortical astrocytes. Science 376, 1441–1446 (2022).

7. You, Z. et al. Mapping of clonal lineages across developmental stages in human neural differentiation. Cell Stem Cell 30, (2023).

8. Guo, Z. et al. RGCC balances self-renewal and neuronal differentiation of neural stem cells in the developing mammalian neocortex. EMBO Rep 22, (2021).

9. Inak, G. et al. Defective metabolic programming impairs early neuronal morphogenesis in neural cultures and an organoid model of Leigh syndrome. Nat Commun 12, (2021).

10. Kang, Y. et al. A human forebrain organoid model of fragile X syndrome exhibits altered neurogenesis and highlights new treatment strategies. Nat Neurosci 24, 1377–1391 (2021).

11. Shimada, I. S., Acar, M., Burgess, R. J., Zhao, Z. & Morrison, S. J. Prdm16 is required for the maintenance of neural stem cells in the postnatal forebrain and their differentiation into ependymal cells. Genes Dev 31, (2017).

12. Shimada, I. S. et al. Derepression of sonic hedgehog signaling upon Gpr161 deletion unravels forebrain and ventricular abnormalities. Dev Biol 450, 47–62 (2019).

13. Mukhopadhyay, S. et al. The ciliary G-protein-coupled receptor Gpr161 negatively regulates the sonic hedgehog pathway via cAMP signaling. Cell 152, 210–223 (2013).

14. Wang, L. et al. Loss of NARS1 impairs progenitor proliferation in cortical brain organoids and leads to microcephaly. Nat Commun 11, (2020).

15. Lancaster, M. A. et al. Cerebral organoids model human brain development and microcephaly. Nature 501, 373–379 (2013).

16. Mill, P., Christensen, S. T. & Pedersen, L. B. Primary cilia as dynamic and diverse signalling hubs in development and disease. Nat Rev Genet 24, 421– 441 (2023).

17. Derderian, C., Canales, G. I. & Reiter, J. F. Seriously cilia: A tiny organelle illuminates evolution, disease, and intercellular communication. Dev Cell 58, 1333–1349 (2023).

18. Casas Gimeno, G. & Paridaen, J. T. M. L. The Symmetry of Neural Stem Cell and Progenitor Divisions in the Vertebrate Brain. Frontiers in Cell and Developmental Biology vol. 10 Preprint at 10.3389/fcell.2022.885269 (2022).

19. Badgandi, H. B., Hwang, S.-H., Shimada, I. S., Loriot, E. & Mukhopadhyay, S. Tubby family proteins are adapters for ciliary trafficking of integral membrane proteins. J Cell Biol 216, (2017).

20. Shimada, I. S. & Kato, Y. Ciliary signaling in stem cells in health and disease: Hedgehog pathway and beyond. Semin Cell Dev Biol 129, 115–125 (2022).

21. Somatilaka, B. N. et al. Ankmy2 Prevents Smoothened-Independent Hyperactivation of the Hedgehog Pathway via Cilia-Regulated Adenylyl Cyclase Signaling. Dev Cell 54, 710–726.e8 (2020).

22. Truong, M. E. et al. Vertebrate cells differentially interpret ciliary and extraciliary cAMP. Cell 184, 2911–2926.e18 (2021).

23. Sheu, S. H. et al. A serotonergic axon-cilium synapse drives nuclear signaling to alter chromatin accessibility. Cell 185, 3390–3407.e18 (2022).

24. Hasenpusch-Theil, K. & Theil, T. The Multifaceted Roles of Primary Cilia in the Development of the Cerebral Cortex. Front Cell Dev Biol 9, 86 (2021).

25. Reiter, J. F. & Leroux, M. R. Genes and molecular pathways underpinning ciliopathies. Nat Rev Mol Cell Biol 18, 533–547 (2017).

26. Zhang, Y. & Beachy, P. A. Cellular and molecular mechanisms of Hedgehog signalling. Nat Rev Mol Cell Biol 24, (2023).

27. Xu, S. & Tang, C. Cholesterol and Hedgehog Signaling: Mutual Regulation and Beyond. Frontiers in Cell and Developmental Biology vol. 10 Preprint at 10.3389/fcell.2022.774291 (2022).

28. Zhang, Y. et al. Structural Basis for Cholesterol Transport-like Activity of the Hedgehog Receptor Patched. Cell 175, (2018).

29. Qi, X. et al. Cryo-EM structure of oxysterol-bound human Smoothened coupled to a heterotrimeric Gi. Nature 571, (2019).

30. Niewiadomski, P. et al. Gli protein activity is controlled by multisite phosphorylation in vertebrate hedgehog signaling. Cell Rep 6, 168–181 (2014).

31. Hoppe, N. et al. GPR161 structure uncovers the redundant role of sterol-regulated ciliary cAMP signaling in the Hedgehog pathway. Nat Struct Mol Biol (2024) doi:10.1038/S41594-024-01223-8.

32. Nie, Y. et al. Specific binding of GPR174 by endogenous lysophosphatidylserine leads to high constitutive Gs signaling. Nat Commun 14, (2023).

33. Pal, K. et al. Smoothened determines β-arrestin-mediated removal of the G protein-coupled receptor Gpr161 from the primary cilium. Journal of Cell Biology 212, (2016).

34. Tang, M., Luo, S. X., Tang, V. & Huang, E. J. Temporal and spatial requirements of Smoothened in ventral midbrain neuronal development. Neural Dev 8, (2013).

35. Gazea, M. et al. Primary cilia are critical for Sonic hedgehog-mediated dopaminergic neurogenesis in the embryonic midbrain. Dev Biol 409, (2016).

36. Willaredt, M. A. et al. A crucial role for primary cilia in cortical morphogenesis. Journal of Neuroscience 28, (2008).

37. Besse, L. et al. Primary cilia control telencephalic patterning and morphogenesis via Gli3 proteolytic processing. Development 138, (2011).

38. Wang, L., Hou, S. & Han, Y. G. Hedgehog signaling promotes basal progenitor expansion and the growth and folding of the neocortex. Nat Neurosci 19, 888– 896 (2016).

39. Matsumoto, N., Tanaka, S., Horiike, T., Shinmyo, Y. & Kawasaki, H. A discrete subtype of neural progenitor crucial for cortical folding in the gyrencephalic mammalian brain. Elife 9, (2020).

40. Van Heurck, R. et al. CROCCP2 acts as a human-specific modifier of cilia dynamics and mTOR signaling to promote expansion of cortical progenitors. Neuron 111, 65–80.e6 (2023).

41. Dessaud, E., McMahon, A. P. & Briscoe, J. Pattern formation in the vertebrate neural tube: A sonic hedgehog morphogen-regulated transcriptional network. Development vol. 135 Preprint at 10.1242/dev.009324 (2008).

42. Pal, K. & Mukhopadhyay, S. Primary cilium and sonic hedgehog signaling during neural tube patterning: Role of GPCRs and second messengers. Dev Neurobiol 75, 337–348 (2015).

43. Nikolopoulou, E., Galea, G. L., Rolo, A., Greene, N. D. E. & Copp, A. J. Neural tube closure: Cellular, molecular and biomechanical mechanisms. Development (Cambridge) vol. 144 Preprint at 10.1242/dev.145904 (2017).

44. Ulloa, F. & Martí, E. Wnt won the war: Antagonistic role of Wnt over Shh controls dorso-ventral patterning of the vertebrate neural tube. Developmental Dynamics vol. 239 Preprint at 10.1002/dvdy.22058 (2010).

45. Le Dréau, G. & Martí, E. Dorsal-ventral patterning of the neural tube: A tale of three signals. Dev Neurobiol 72, (2012).

46. Jacob, J. & Briscoe, J. Gli proteins and the control of spinal-cord patterning. EMBO Rep 4, 761–765 (2003).

47. Dias, J. M. et al. A Shh/Gli-driven three-node timer motif controls temporal identity and fate of neural stem cells. Sci Adv 6, (2020).

48. Delás, M. J. et al. Developmental cell fate choice in neural tube progenitors employs two distinct cis-regulatory strategies. Dev Cell 58, (2023).

49. Bhaduri, A. et al. An atlas of cortical arealization identifies dynamic molecular signatures. Nature 598, 200–204 (2021).

50. Uzquiano, A. et al. Proper acquisition of cell class identity in organoids allows definition of fate specification programs of the human cerebral cortex. Cell 185, 3770–3788.e27 (2022).

51. Eiraku, M. et al. Self-Organized Formation of Polarized Cortical Tissues from ESCs and Its Active Manipulation by Extrinsic Signals. Cell Stem Cell 3, (2008).

52. Bowles, K. R. et al. ELAVL4, splicing, and glutamatergic dysfunction precede neuron loss in MAPT mutation cerebral organoids. Cell 184, 4547–4563.e17 (2021).

53. Rossi, G., Manfrin, A. & Lutolf, M. P. Progress and potential in organoid research. Nat Rev Genet 19, 671–687 (2018).

54. Hofer, M. & Lutolf, M. P. Engineering organoids. Nat Rev Mater 6, 402–420 (2021).

55. Sakaguchi, H. et al. Generation of functional hippocampal neurons from self-organizing human embryonic stem cell-derived dorsomedial telencephalic tissue. Nat Commun 6, (2015).

56. Suga, H. et al. Self-formation of functional adenohypophysis in three-dimensional culture. Nature 480, (2011).

57. Schembs, L. et al. The ciliary gene INPP5E confers dorsal telencephalic identity to human cortical organoids by negatively regulating Sonic hedgehog signaling. Cell Rep 39, (2022).

58. Gabriel, E. et al. CPAP promotes timely cilium disassembly to maintain neural progenitor pool. EMBO J 35, 803–819 (2016).

59. Sloan, S. A., Andersen, J., PaLca, A. M., Birey, F. & PaLca, S. P. Generation and assembly of human brain region-specific three-dimensional cultures. Nat Protoc 13, 2062–2085 (2018).

60. Higginbotham, H. et al. Arl13b-regulated cilia activities are essential for polarized radial glial scaffold formation. Nat Neurosci 16, 1000–1007 (2013).

61. Bay, S. N., Long, A. B. & Caspary, T. Disruption of the ciliary GTPase Arl13b suppresses Sonic hedgehog overactivation and inhibits medulloblastoma formation. Proc Natl Acad Sci U S A 115, 1570–1575 (2018).

62. Caspary, T., Larkins, C. E. & Anderson, K. V. The Graded Response to Sonic Hedgehog Depends on Cilia Architecture. Dev Cell 12, 767–778 (2007).

63. Liem, K. F. et al. The IFT-A complex regulates Shh signaling through cilia structure and membrane protein trafficking. Journal of Cell Biology 197, (2012).

64. Wang, C. et al. Centrosomal protein Dzip1l binds Cby, promotes ciliary bud formation, and acts redundantly with bromi to regulate ciliogenesis in the mouse. Development (Cambridge) 145, (2018).

65. Ko, H. W. et al. Broad-Minded Links Cell Cycle-Related Kinase to Cilia Assembly and Hedgehog Signal Transduction. Dev Cell 18, (2010).

66. Horner, V. L. & Caspary, T. Disrupted dorsal neural tube BMP signaling in the cilia mutant Arl13bhnn stems from abnormal Shh signaling. Dev Biol 355, (2011).

67. Cohen, M. et al. Ptch1 and Gli regulate Shh signalling dynamics via multiple mechanisms. Nat Commun 6, (2015).

68. Kutejova, E., Sasai, N., Shah, A., Gouti, M. & Briscoe, J. Neural Progenitors Adopt Specific Identities by Directly Repressing All Alternative Progenitor Transcriptional Programs. Dev Cell 36, 639–653 (2016).

69. Shimada, I. S. et al. Basal Suppression of the Sonic Hedgehog Pathway by the G-Protein-Coupled Receptor Gpr161 Restricts Medulloblastoma Pathogenesis. Cell Rep 22, (2018).

70. Pusapati, G. V. et al. G protein-coupled receptors control the sensitivity of cells to the morphogen Sonic Hedgehog. Sci Signal 11, (2018).

71. Nozaki, S. et al. Regulation of ciliary retrograde protein trafficking by the Joubert syndrome proteins ARL13B and INPP5E. J Cell Sci 130, 563–576 (2017).

72. Qiu, H., Fujisawa, S., Nozaki, S., Katoh, Y. & Nakayama, K. Interaction of INPP5E with ARL13B is essential for its ciliary membrane retention but dispensable for its ciliary entry. Biol Open 10, (2021).

73. Hwang, S. H. et al. The G protein-coupled receptor Gpr161 regulates forelimb formation, limb patterning and skeletal morphogenesis in a primary cilium-dependent manner. Development (Cambridge) 145, (2018).

74. Kim, S. E. et al. Dominant negative GPR161 rare variants are risk factors of human spina bifida. Hum Mol Genet 28, (2019).

75. Fleck, J. S. et al. Inferring and perturbing cell fate regulomes in human brain organoids. Nature 621, 365–372 (2023).

76. Moore, B. S. et al. Cilia have high cAMP levels that are inhibited by Sonic Hedgehog-regulated calcium dynamics. Proc Natl Acad Sci U S A 113, 13069– 13074 (2016).

77. Jiang, J. Y., Falcone, J. L., Curci, S. & Hofer, A. M. Direct visualization of cAMP signaling in primary cilia reveals up-regulation of ciliary GPCR activity following Hedgehog activation. Proc Natl Acad Sci U S A 116, 12066–12071 (2019).

78. Hwang, S. H., Somatilaka, B. N., White, K. & Mukhopadhyay, S. Ciliary and extraciliary gpr161 pools repress hedgehog signaling in a tissue-specific manner. Elife 10, (2021).

79. Kim, S.-E. et al. Wnt1 Lineage Specific Deletion of Gpr161 Results in Embryonic Midbrain Malformation and Failure of Craniofacial Skeletal Development. Front Genet 0, 2366 (2021).

80. Hwang, S. H., White, K. A., Somatilaka, B. N., Wang, B. & Mukhopadhyay, S. Context-dependent ciliary regulation of hedgehog pathway repression in tissue morphogenesis. PLoS Genet 19, (2023).

81. Legué, E. & Liem, K. F. Mutations in Ciliary Trafficking Genes affect Sonic Hedgehog-dependent Neural Tube Patterning Differentially along the Anterior– Posterior Axis. Neuroscience 450, 3–14 (2020).

82. Brooks, E. R., Islam, M. T., Anderson, K. V. & Zallen, J. A. Sonic hedgehog signaling directs patterned cell remodeling during cranial neural tube closure. Elife 9, 1–36 (2020).

83. May, E. A., Sroka, T. J. & Mick, D. U. Phosphorylation and Ubiquitylation Regulate Protein Trafficking, Signaling, and the Biogenesis of Primary Cilia. Front Cell Dev Biol 9, (2021).

84. Garcia-Gonzalo, F. R. et al. Phosphoinositides Regulate Ciliary Protein Trafficking to Modulate Hedgehog Signaling. Dev Cell 34, 400–409 (2015).

85. Schou, K. B., Pedersen, L. B. & Christensen, S. T. Ins and outs of GPCR signaling in primary cilia. EMBO Rep 16, 1099–1113 (2015).

86. Gigante, E. D., Taylor, M. R., Ivanova, A. A., Kahn, R. A. & Caspary, T. Arl13b regulates sonic hedgehog signaling from outside primary cilia. Elife 9, (2020).

87. Suciu, S. K., Long, A. B. & Caspary, T. Smoothened and ARL13B are critical in mouse for superior cerebellar peduncle targeting. Genetics 218, (2021).

88. Su, C. Y., Bay, S. N., Mariani, L. E., Hillman, M. J. & Caspary, T. Temporal deletion of Arl13b reveals that a mispatterned neural tube corrects cell fate over time. Development (Cambridge) 139, (2012).

89. Fiore, L. et al. Optic vesicle morphogenesis requires primary cilia. Dev Biol 462, (2020).

90. Larkins, C. E., Gonzalez Aviles, G. D., East, M. P., Kahn, R. A. & Caspary, T. Arl13b regulates ciliogenesis and the dynamic localization of Shh signaling proteins. Mol Biol Cell 22, (2011).

91. Kumamoto, N. et al. A role for primary cilia in glutamatergic synaptic integration of adult-born neurons. Nat Neurosci 15, (2012).

92. Breunig, J. J. et al. Primary cilia regulate hippocampal neurogenesis by mediating sonic hedgehog signaling. Proc Natl Acad Sci U S A 105, (2008).

93. Das, R. M. & Storey, K. G. Apical abscission alters cell polarity and dismantles the primary cilium during neurogenesis. Science (1979) 343, 200–204 (2014).

94. Kondo, Y. et al. Histone deacetylase inhibitor valproic acid promotes the differentiation of human induced pluripotent stem cells into hepatocyte-like cells. PLoS One 9, (2014).

95. Iwao, T. et al. Differentiation of human induced pluripotent stem cells into functional enterocyte-like cells using a simple method. Drug Metab Pharmacokinet 29, 44–51 (2014).

96. Nishino, K. et al. Defining hypo-methylated regions of stem cell-specific promoters in human iPS cells derived from extra-embryonic amnions and lung fibroblasts. PLoS One 5, (2010).

97. Nakamura, Y. et al. Biallelic null variants in PNPLA8 cause microcephaly through the reduced abundance of basal radial glia. medRxiv (2023) doi:10.1101/2023.04.26.23288947.

98. Miyazaki, T., Isobe, T., Nakatsuji, N. & Suemori, H. Efficient Adhesion Culture of Human Pluripotent Stem Cells Using Laminin Fragments in an Uncoated Manner. Sci Rep 7, (2017).

99. Bak, R. O. & Porteus, M. H. CRISPR-Mediated Integration of Large Gene Cassettes Using AAV Donor Vectors. Cell Rep 20, 750–756 (2017).

100. Oceguera-Yanez, F. et al. Engineering the AAVS1 locus for consistent and scalable transgene expression in human iPSCs and their differentiated derivatives. Methods 101, (2016).

101. Golden, R. J. et al. An Argonaute phosphorylation cycle promotes microRNA-mediated silencing. Nature 542, (2017).

102. Xiang, Y. et al. hESC-Derived Thalamic Organoids Form Reciprocal Projections When Fused with Cortical Organoids. Cell Stem Cell 24, 487–497.e7 (2019).

103. Meharena, H. S. et al. Down-syndrome-induced senescence disrupts the nuclear architecture of neural progenitors. Cell Stem Cell 29, (2022).

104. Zhang, W. et al. Cerebral organoid and mouse models reveal a RAB39b-PI3K-mTOR pathway-dependent dysregulation of cortical development leading to macrocephaly/ autism phenotypes. Genes Dev 34, (2020).

105. Hanaichi, T. et al. A Stable Lead by Modification of Sato’s Method. J Electron Microsc (Tokyo) 35, 304–306 (1986).

106. Chen, S., Zhou, Y., Chen, Y. & Gu, J. Fastp: An ultra-fast all-in-one FASTQ preprocessor. in Bioinformatics vol. 34 (2018).

107. Langmead, B. & Salzberg, S. L. Fast gapped-read alignment with Bowtie 2. Nat Methods 9, (2012).

108. Danecek, P. et al. Twelve years of SAMtools and BCFtools. Gigascience 10, (2021).

109. Ramírez, F. et al. deepTools2: a next generation web server for deep-sequencing data analysis. Nucleic Acids Res 44, (2016).

110. Heinz, S. et al. Simple Combinations of Lineage-Determining Transcription Factors Prime cis-Regulatory Elements Required for Macrophage and B Cell Identities. Mol Cell 38, (2010).

111. Robinson, J. T. et al. Integrative genomics viewer. Nature Biotechnology vol. 29 Preprint at 10.1038/nbt.1754 (2011).

